# Learning a genome-wide score of human-mouse conservation at the functional genomics level

**DOI:** 10.1101/2020.09.08.288092

**Authors:** Soo Bin Kwon, Jason Ernst

## Abstract

Identifying genomic regions with functional genomic properties that are conserved between human and mouse is an important challenge in the context of mouse model studies. To address this, we take a novel approach and learn a score of evidence of conservation at the functional genomics level by integrating large-scale information in a compendium of epigenomic, transcription factor binding, and transcriptomic data from human and mouse. The computational method we developed to do this, Learning Evidence of Conservation from Integrated Functional genomic annotations (LECIF), trains a neural network, which is then used to generate a genome-wide score in human and mouse. The resulting LECIF score highlights human and mouse regions with shared functional genomic properties and captures correspondence of biologically similar human and mouse annotations even though it was not explicitly given such information. LECIF will be a resource for mouse model studies.

## Introduction

Genome-wide maps of chromatin accessibility, transcription factor binding, histone modifications, and gene expression data across diverse cell and tissue types^1–4^ have been resources for interpreting disease associated genetic variation^5–10^. Many studies interrogate human loci of interest by perturbing their homologous loci in mouse^11–17^. A key question in doing so is the extent to which the homologous loci in mouse have similar roles to the human loci. Conversely, loci associated with phenotypes can be discovered in mouse first, raising the question of the degree to which their properties are shared with human^18^.

A relatively large percentage of the human genome, 40%, has a homologous locus in the mouse genome as determined by pairwise sequence alignment^19^. However, a much smaller fraction of these aligning pairs of loci are constrained at the sequence level^20–23^. It is unclear to what extent homologous loci in human and mouse lacking sequence constraint still share other properties, in particular, functional genomic properties. With large-scale functional genomic resources that have become available in mouse^24,25^ in addition to human, there is an opportunity to more systematically and confidently detect conservation at the functional genomics level between these species.

Previous work comparing cross-species functional genomics data to infer conservation have largely focused on comparing pairs of matched experiments for the same assay in a corresponding cell or tissue type across species^26–36^. While useful, comparing a pair of experiments from two species can be underpowered to differentiate evidence of conservation from similarity observed by chance. Studies that jointly compare multiple pairs of experiments instead are able to consider information from multiple biological conditions and potentially gain power in inferring conservation of functional genomic properties^27,33,35,36^. However, such approaches have often relied on manually matching corresponding experiments and have not been scaled to leverage the vast amounts of diverse of data available in both human and mouse. The challenge in taking advantage of such data is that many experiments do not have an obvious corresponding experiment, and even when one is assumed there could in practice be confounding differences. Previous work partly addressed some of these issues^24,37–43^, but still limited their work to either one data type or dimension of the data at a time and thus only utilized a small fraction of the available data to find evidence of conservation.

Given the increasingly diverse functional genomic resources available for human and mouse, there is a need for an integrative method that can better leverage those resources to infer conservation at the functional genomics level and identify regions with functional genomic properties shared between human and mouse. Thus, here we developed Learning Evidence of Conservation from Integrated Functional genomic annotations (LECIF), a supervised machine learning approach that quantifies evidence of conservation based on large-scale functional genomic data from a pair of species, which we applied to human and mouse. While LECIF takes advantage of data from a diverse set of cell types collected by various assays, it does not require explicit matching of experiments from different species by their biological source or data type. LECIF uses pairwise sequence alignment data only to label training examples, inferring conservation specifically from functional genomics data and not from DNA sequence. We applied LECIF to a compendium of thousands of human and mouse functional genomic annotations and learned a human-mouse LECIF score for every pair of human and mouse genomic regions that align at the sequence level.

We conducted various analyses to understand the properties of the LECIF score as well as the information it provides for variant interpretation and translation of mouse model studies. The LECIF score is highly predictive of pairs of regions that align at the sequence level, even when controlling for properties of regions that align in general. The score captures correspondence of biologically similar annotations between human and mouse, even though LECIF was not explicitly given such information. While the LECIF score is moderately correlated with sequence constraint scores, it captures distinct information on conserved properties. Regions with high LECIF score show enrichment for phenotypic associated genetic variants, even when conditioned on sequence constrained elements. The LECIF score is preferentially higher in regions previously shown to have similar phenotypic properties in human and mouse at the genetic and epigenetic level. We expect the human-mouse LECIF score will be an important a resource for studies using mouse as a model organism.

## Results

### Learning evidence of conservation from integrated functional genomic annotations

LECIF quantifies evidence of conservation between genomic regions from two different species at the functional genomics level based on a large and diverse set of functional genomic annotations (**Fig. 1**). LECIF uses functional genomic features as input to an ensemble of 100 neural networks where sequence alignment information is used to label training data, but not as features (**Methods**). For training data, positive examples are pairs of regions between the two species that align at the sequence level while negative examples are randomly mismatched pairs of human and mouse regions that align somewhere in the other species, but not to each other (**Fig. 1a**). LECIF assumes that positive examples are more likely to be conserved at the functional genomics level than negative examples. Since all human and mouse regions in the negative examples still align somewhere in the other genome, this allows LECIF to learn pairwise characteristics of aligning human and mouse regions instead of learning the characteristics of regions that align to the other genome in general. We provided the neural network classifier with more than two million positive and two million negative training examples.

**Figure 1.**
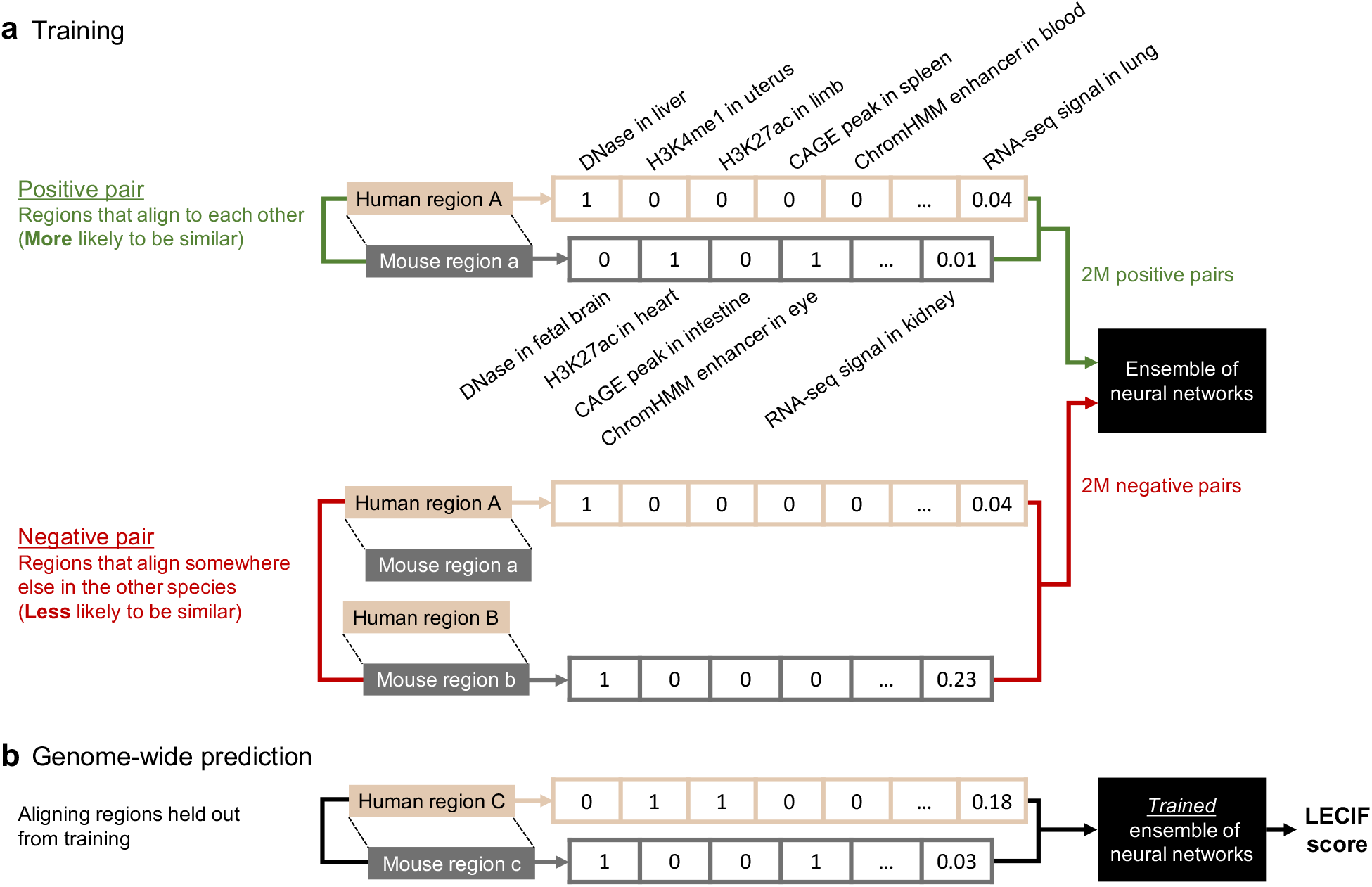
Overview of the LECIF method. **a.** Supervised learning procedure of LECIF. For every pair of human and mouse genomic regions, two feature vectors are generated from their functional genomic annoations, one vector for the human region (beige) and the other vector for the mouse region (gray). Each feature vector consists of thousands of functional genomic annotations, as listed in **Supplementary Table 1.** Only a subset of the features are shown here. These two species-specific feature vectors are given to an ensemble of neural networks (ENN), which is trained to distinguish positive pairs (green), which are aligning human and mouse regions, from negative pairs (red), which are randomly mismatched human and mouse regions that do not align to each other, but somewhere else in the other species. Here we provide about two million positive and two million negative training examples. Feature labels (e.g. DNase in liver) and matching of features across species are not provided to LECIF. **b.** Genome-wide prediction procedure of LECIF. Once trained as illustrated in **a**, the ENN can estimate the probability of any given pair of human and mouse regions being classified as a positive pair. We consider this probability, the LECIF score, to represent the evidence of conservation observed in the functional genomics data annotating the given pair. Here we generate the LECIF score for all pairs of aligning human and mouse regions. When generating a prediction for a pair, LECIF uses an ENN trained on data excluding the pair as described in **Methods** and **Supplementary Table 2**.

For each example there were more than 8,000 human and 3,000 mouse functional genomic features defined (**Fig. 2a**). Among these features were binary features corresponding to whether a genomic base overlapped with peak calls from DNase-seq experiments, ChIP-seq experiments of transcription factors (TF), histone modifications and histone variants, and Cap Analysis of Gene Expression (CAGE) experiments. Additionally, there were binary features corresponding to each state and tissue type of ChromHMM^44^ chromatin state annotations and numerical features corresponding to normalized signals from RNA-seq experiments. These data covered a wide range of cell and tissue types and were generated by the ENCODE^2^, Mouse ENCODE^24^, Roadmap Epigenomics Project^3^, or FANTOM5^45^ consortia (**Methods**; **Supplementary Table 1**). We did not provide pairwise alignment or DNA sequence information as features to the classifier so that LECIF infers conservation specifically at the functional genomics level rather than at the sequence level.

**Figure 2.**
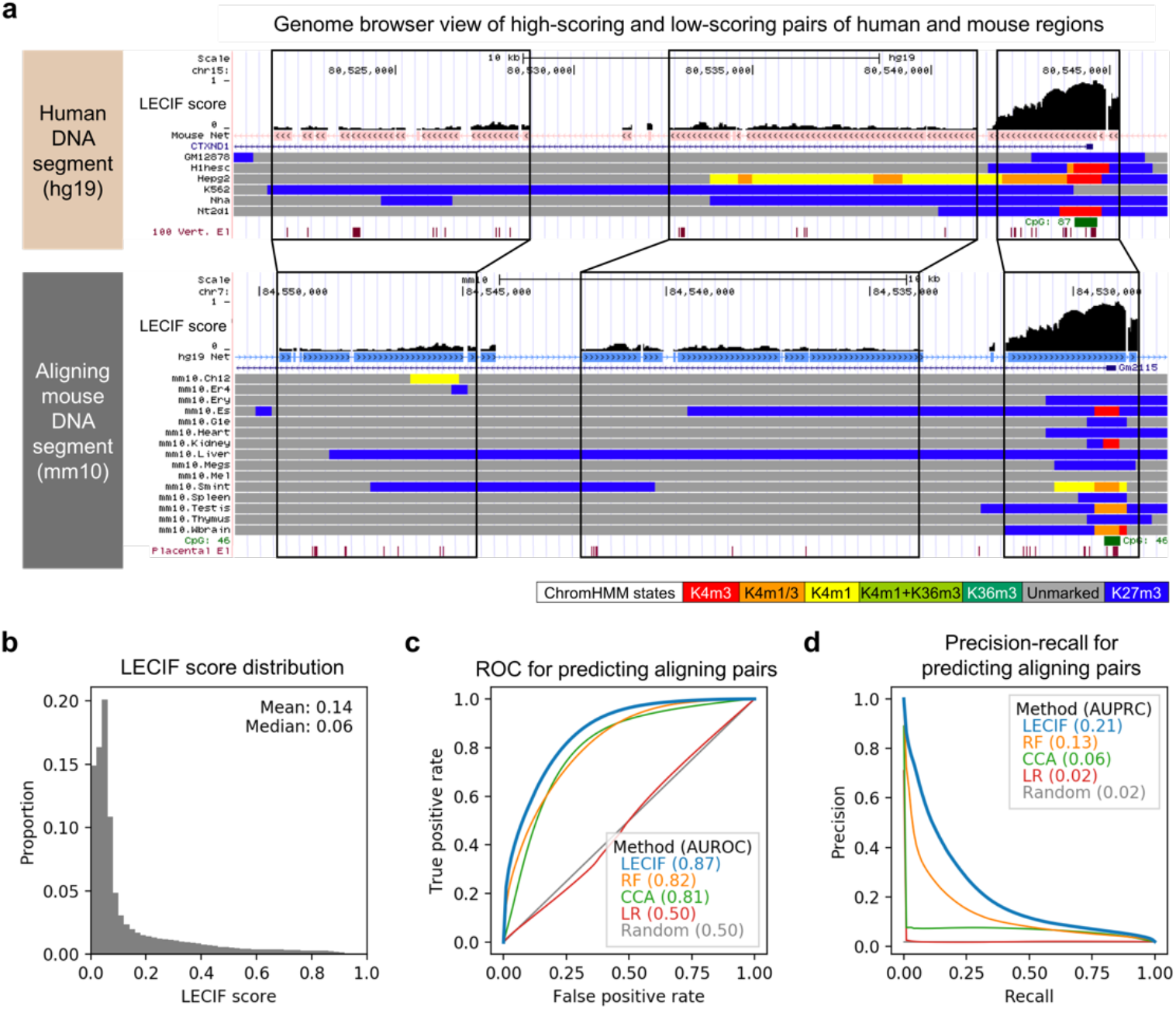
Characteristics of the LECIF score. **a.** Genome Browser^46^ views illustrating the human-mouse LECIF score annotating human protein-coding gene CTXND1 (top) and its mouse ortholog Gm2115 (bottom). In each browser view, LECIF score is shown in the top track in black. Below the score, the net pairwise sequence alignment annotation^19^ marks regions that align between human and mouse indicated with salmon and blue boxes, respectively. Regions not annotated with a colored box in the net track have no human-mouse sequence alignment and thus no LECIF score. Below the net track is the RefSeq gene annotation. The next set of tracks below that are ChromHMM chromatin state annotations^44^ for different cell and tissue types, from a chromatin state model that was previously learned jointly for human and mouse^24^. Legend with the chromatin state labels as previously defined^24^ is shown in the bottom right. Below the chromatin state annotations are CpG island and PhastCons element^20^ annotations. Black solid lines highlight DNA segments that largely align to each other. The mouse genome browser view is shown in the reverse direction (3’ to 5’) because that is the direction in which mouse aligns to human in this DNA segment. **b.** Distribution of the LECIF score, which is right-skewed between 0 and 1 with a genome-wide mean of 0.14 and median of 0.06. Fifty equal-width bins were used to plot the histogram. **c.** Receiver operating characteristic (ROC) curve comparing LECIF, random forest (RF), canonical correlation analysis (CCA), and logistic regression (LR) for classifying pairs of regions that align at the sequence level, evaluated on a common set of held-out test data. Shown on the y=x line is random expectation. Legend indicates color and mean area under the ROC curve (AUROC) corresponding to each method. The ROC curve and AUROC of each method was obtained by classifying 100,000 positive and 100,000 negative examples randomly sampled with replacement from all available test examples 100 times. Negative examples were weighted 50 times more than positive examples. For each method, standard deviation of the 100 AUROC values was under 0.005. **d.** Similar analysis as in **c** except for precision-recall (PR) curves instead of ROC curves. Standard deviation of the 100 area under the PR curve (AUPRC) values was also under 0.005 for all methods.

After training, we used the trained classifier to make genome-wide predictions, annotating both the human and mouse genomes with the human-mouse LECIF score **(Fig. 1b**). We designed the LECIF score to highlight regions with strong evidence of conservation at the functional genomics level by weighting negative examples 50 times more than positive examples during training. We confirmed that the relative ranking of predictions was robust to different weightings (**Supplementary Fig. 1**). We made predictions at every 50 bp within each alignment block as it was computationally more efficient than doing so at every base, and we confirmed that this score was strongly correlated with a score computed at a single base resolution with a Pearson correlation coefficient (PCC) of 0.99 (**Methods**). Additionally, we confirmed that the LECIF score had strong agreement when trained on non-overlapping sets of chromosomes (PCC: 0.90; **Methods**) and that ensembling 100 neural networks led to better agreement than using a single neural network (**Supplementary Fig. 2**).

### LECIF outperforms baseline methods in predicting aligning pairs

We evaluated LECIF and baseline methods at predicting whether pairs of regions that were held out from training align at the sequence level. LECIF had strong predictive power for this with an area under the receiver operating characteristic curve (AUROC) of 0.87 and an area under the precision-recall curve (AUPRC) of 0.23 compared to a random expectation of 0.50 and 0.02, respectively (**Fig. 2c,d**).

We compared LECIF to random forest (RF), canonical correlation analysis (CCA), and logistic regression (LR) (**Fig. 2c,d**). When classifying held-out test examples, LECIF outperformed RF and CCA, with statistically significantly better AUROC and AUPRC (RF AUROC: 0.82; CCA AUROC: 0.81; RF AUPRC: 0.13; CCA AUPRC: 0.06; Wilcoxon signed-rank test *P*<0.0001). As expected, logistic regression had no predictive power (AUROC: 0.50; AUPRC: 0.02), since it only considers features marginally and the positive and negative examples were defined such that each feature has an identical marginal distribution in positive and negative data.

### Assessment of predictive power for aligning pairs of human regions and extended mouse regions

The LECIF method is not limited to scoring pairs of human and mouse regions that align at the sequence level. Previous comparative studies have reported movements of regulatory elements during evolution, where homologous regulatory activity of a human region is found in a region near the aligning region in another species instead of the aligning region^26,47,48^. This could suggest that for a given human region, instead of focusing on the LECIF score specifically at the aligning mouse region, it could be advantageous to consider also the scores at non-aligning mouse regions that are proximal to the mouse region aligning to human. However, such an approach also increases the possibility of high-scoring pairs that are false positives.

We thus investigated an approach where for a given aligning pair of human and mouse regions we took the maximum LECIF score from pairs consisting of the human region and any mouse region located within a window centered around the aligning mouse region (**Methods**; **Supplementary Fig. 3**). We varied window sizes and repeated the same AUC evaluations for predicting aligning regions as above (**Supplementary Fig. 4**). We found that as we expanded the window size the predictive power decreased. When we trained LECIF with an alternative set of negative examples selected from a genome background and repeated the evaluations (**Methods**), the expanded window still had decreased predictive power (**Supplementary Fig. 4**). These results motivated us to focus our application of LECIF to aligning regions. We note that because of the resolution at which the LECIF score is defined, the score may still be capturing small movements of regulatory sites even without explicitly expanding the window.

### Distribution of LECIF score for different input features

We next investigated the distribution of the LECIF score overlapping the functional genomic annotations provided to LECIF as binary input features. For each DNase-seq, ChIP-seq, or CAGE feature, we computed the mean LECIF score of the annotated regions. We saw that regions with signatures of regulatory activity had a mean LECIF score above the genome-wide mean (**Fig. 3a**). On average, human regions annotated within a peak call from a DNase-seq, TF ChIP-seq, or CAGE experiment had mean LECIF scores of 0.39, 0.47, and 0.70, respectively, compared to the genome-wide mean of 0.14. Human regions in peak calls of ChIP-seq for the histone modifications H3K4me3, associated with promoters, and H3K27ac, associated with active enhancers and promoters, had mean LECIF scores of 0.60 and 0.41, respectively. We observed similar trends with annotated bases in the mouse genome (**Supplementary Fig. 5**).

**Figure 3.**
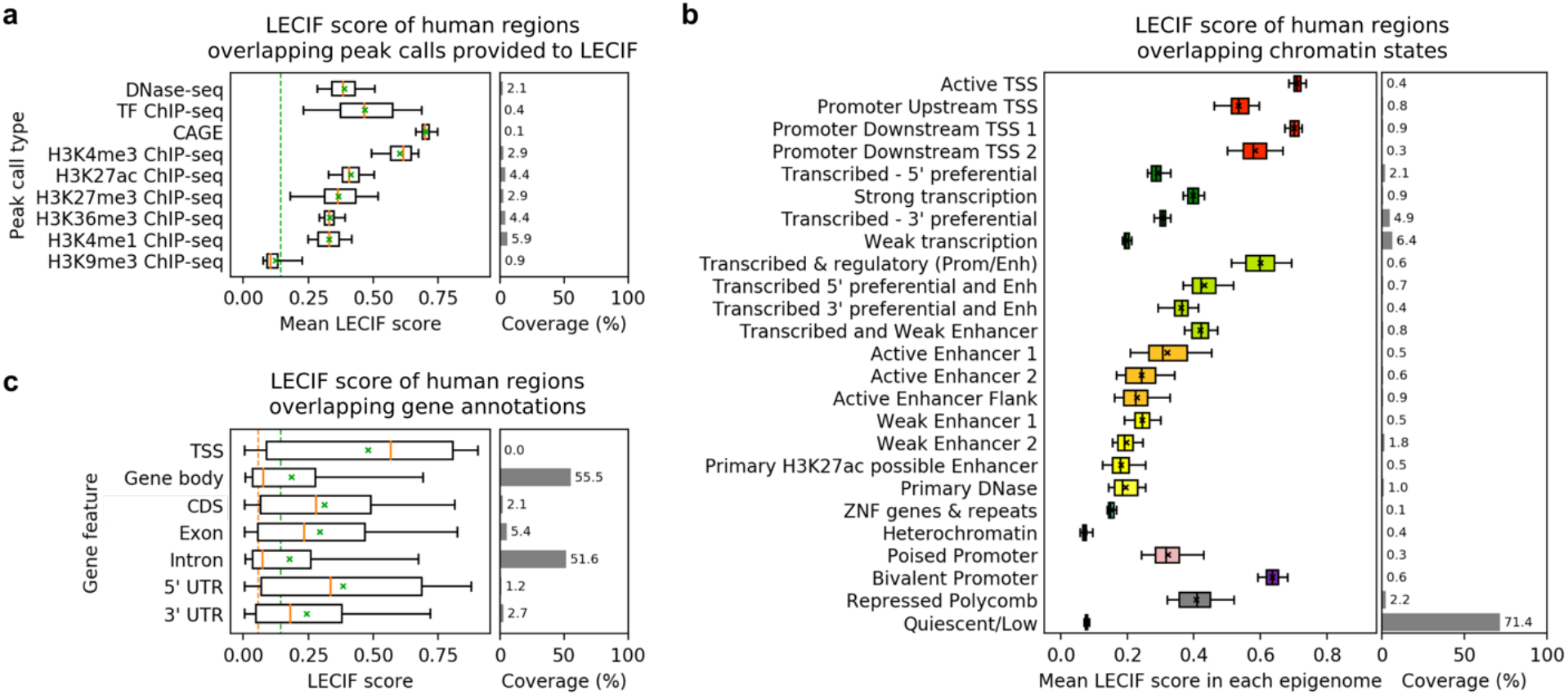
Relationship of LECIF score to functional genomic and gene annotations. **a.** Shown for each type of functional genomic experiments listed is the distribution of mean LECIF score over experiments of that type in human. The mean LECIF score for an experiment is computed based on averaging the LECIF score of regions overlapping a peak call from the experiment. The set of experiments are the same as provided to LECIF as input features. Each distribution is represented by a boxplot with median (orange solid line), mean (green ‘x’), 25^th^ and 75^th^ percentiles (box), and 5^th^ and 95^th^ percentiles (whisker). Green dashed vertical line across the entire panel denotes the genome-wide mean LECIF score. Right panel shows mean coverage of each type of peak call across all human regions that align to the mouse genome. A version of this plot for mouse is shown in **Supplementary Fig. 5**. **b.** Shown for each chromatin state^3,49^ from a model learned in human is the distribution of mean LECIF score over different epigenomes. The mean LECIF score for a chromatin state in an epigenome is computed based on averaging the LECIF score of regions overlapping the chromatin state in the epigenome. Each distribution is represented by a boxplot with median (black vertical line), mean (black ‘x’), 25^th^ and 75^th^ percentiles (box), and 5^th^ and 95^th^ percentiles (whisker). Right panel shows mean coverage of each state across all human regions that align to the mouse genome. A version of this plot for mouse is shown in **Supplementary Fig. 6**. **c.** Distribution of LECIF score in human regions overlapping indicated GENCODE gene feature annotations. Each distribution is represented by a boxplot with median (orange solid line), mean (green ‘x’), 25^th^ and 75^th^ percentiles (box), and 5^th^ and 95^th^ percentiles (whisker). Dashed vertical lines in orange and green across the entire panel denote the genome-wide median and mean LECIF scores, respectively. Right panel shows coverage of each annotation across all human regions that align to the mouse genome. TSS: transcription start site; CDS: coding sequence; UTR: untranslated region. A version of this plot for mouse is shown in **Supplementary Fig. 7**.

We also computed the mean LECIF score for each chromatin state by taking the average of the mean LECIF score for the chromatin state in each epigenome^3^ (**Fig. 3b**). We saw a wide range of mean LECIF scores with the highest mean of 0.71 for an active transcription start site (TSS) state and the lowest means of 0.07 and 0.08 for the heterochromatin and quiescent states, respectively. The relatively high mean we observed for promoter states (0.53 to 0.71) was consistent with a relatively high score for GENCODE TSS annotations compared to other GENCODE annotated gene features, which were not directly provided as input features to LECIF (**Fig. 3c**). Candidate enhancer states outside of transcribed regions had an intermediate mean LECIF score ranging from 0.18 to 0.32, which was lower than the mean scores of promoter associated states and consistent with previous findings that enhancers tend to evolve faster than promoters^29^. We observed these trends with mouse chromatin states and annotated gene features as well (**Supplementary Figs. 6-7**).

### LECIF score highlights human and mouse regions with shared regulatory or transcriptional activity

We next investigated the LECIF score in relation to human and mouse genomic annotations jointly. We first matched a subset of human and mouse ChIP-seq experiments of H3K27ac by their tissue of origin for 14 tissue type groups. We then quantified the cross-species similarity of the peak calls for each pair of regions jointly across the 14 tissue type groups using a weighted Jaccard similarity coefficient (**Methods**). We saw that the LECIF score was positively correlated with the weighted Jaccard similarity coefficient (PCC: 0.45; **Fig. 4a**). This is despite LECIF not being given any information regarding tissue of origin of the experiments in the compendium of functional genomic annotations.

**Figure 4.**
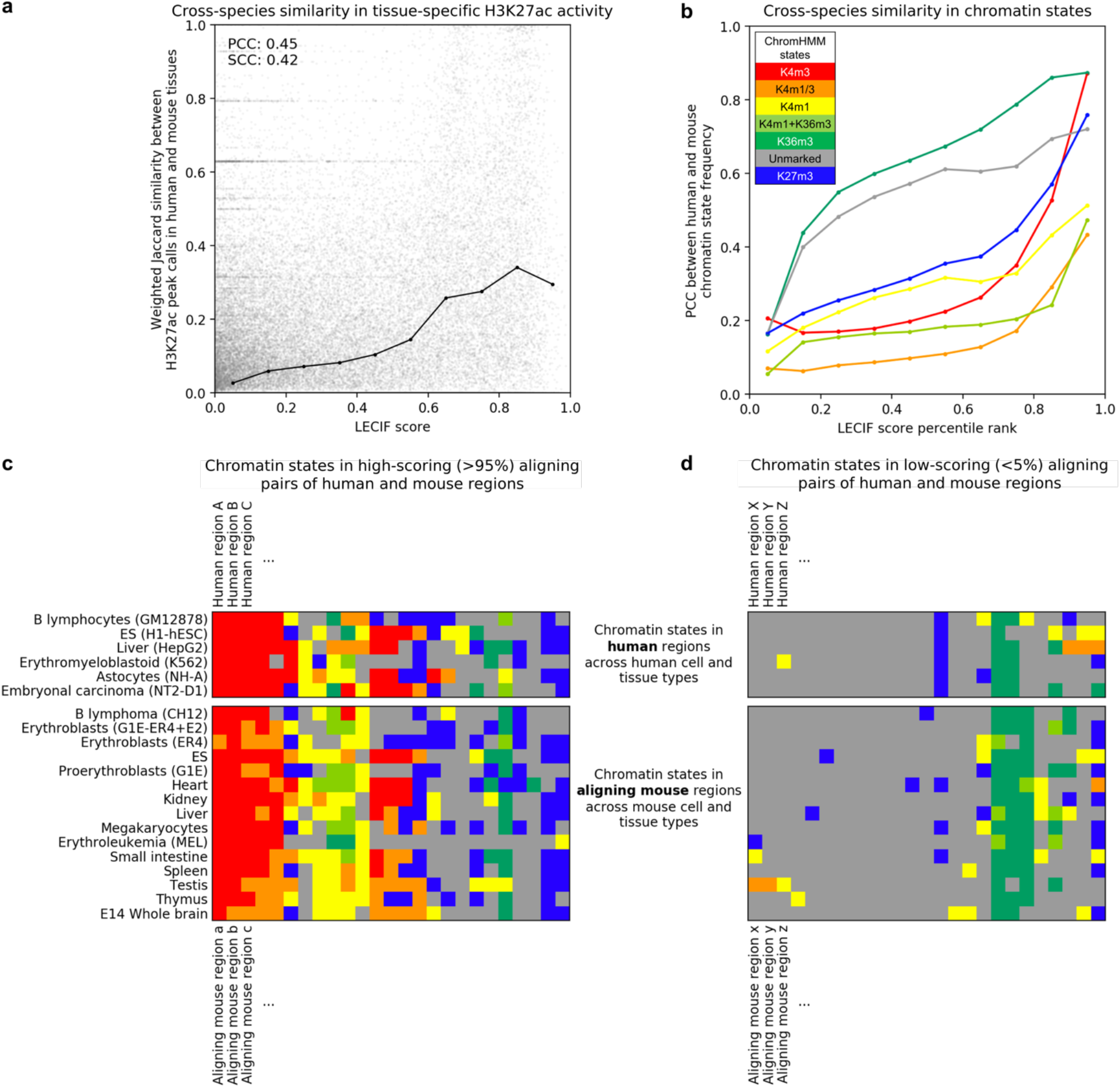
Correspondence of LECIF score to matched human and mouse annotations. **a.** Scatter plot showing with a gray dot for each aligning pair of human and mouse regions the LECIF score (x-axis) and cross-species similarity of matched tissue-specific H3K27ac activity (y-axis). The H3K27ac activity for a region in a tissue type group and species is quantified as the fraction of experiments in the tissue type of the species with peak calls overlapping the region. The cross-species similarity of the H3K27ac activity is quantified as the weighted Jaccard similarity coefficient over 14 matched tissue types (**Methods**). The black circles show the mean similarity coefficient of pairs binned by the LECIF score for ten equal-width bins. These circles are connected with lines based on piecewise linear interpolation. One hundred thousand aligning pairs were randomly selected to plot this scatter plot. A version of this figure for the human-only baseline score is shown in **Supplementary Fig. 10**. **b.** Cross-species agreement in chromatin state^24,44^ frequency in pairs of aligning human and mouse regions binned by the percentile rank of the LECIF score for a ChromHMM model learned jointly between human and mouse. Pairs were binned by their LECIF score percentile rank using ten bins with similar number of pairs in each bin. For each state and aligning regions, we computed the frequency of the state across cell and tissue types for human and mouse separately. We then, for each state and LECIF score percentile rank bin, computed the PCC between the corresponding human and mouse frequencies for that state across all aligning pairs within the same percentile rank bin (**Methods**). The values are shown with colored circles according to the chromatin state legend on the top left from Ref. 23. The circles for a state are connected with lines based on piecewise linear interpolation. Alternative versions of this plot with the pairs binned with a different number of bins and based on the LECIF score instead of its percentile rank are shown in **Supplementary Fig. 8**. **c.** ChromHMM chromatin state^24,44^ annotations in randomly selected pairs of aligning human and mouse regions with high LECIF score. Each row in the top sub-panel corresponds to a human cell or tissue type. Each row in the bottom sub-panel corresponds to a mouse cell or tissue type. Each column is a randomly selected pair of regions with high LECIF score (>95^th^ percentile). Each cell shows the color of the chromatin state with which the human or mouse region (column) is annotated in a specific cell or tissue type (row). The chromatin state model and state coloring are the same as in **b**. Pairs (columns) were ordered based on hierarchical clustering applied to their chromatin state annotations using Ward’s linkage with optimal leaf ordering^50^. **d.** Same as **c,** but with randomly selected pairs with low LECIF score (<5^th^ percentile).

We next examined the LECIF score in relation to the chromatin state annotations of pairs of human and mouse regions. We used the state mappings from a concatenated model of ChromHMM^44^ where a shared set of states were learned for human and mouse^24^. For different ranges of the LECIF score, we correlated the chromatin state frequency between human and mouse across regions in that score range (**Methods**). High-scoring pairs of regions tended to be annotated with similar sets of states in human and mouse epigenomes (**Fig. 4b,c, Supplementary Fig. 8**). In contrast, low-scoring pairs of regions were annotated with dissimilar sets of states in human and mouse and more frequently annotated with the quiescent state than high-scoring pairs (**Fig. 4b,d, Supplementary Figs. 8-9**).

For these analyses, we also verified the advantage of integrating human and mouse data by generating a human-only baseline score, which was defined within human regions that align to mouse using only functional genomics data from human (**Methods**). The score was weakly correlated with the human-mouse LECIF score with a PCC of 0.13. For analyzing H3K27ac peak calls jointly across matched tissue types, the human-only baseline score was not as strongly correlated with the weighted Jaccard similarity coefficient as the LECIF score was (LECIF PCC: 0.45; human-only baseline PCC: 0.08; **Supplementary Fig. 10**). For the analysis of chromatin state assignments, pairs with high human-only baseline scores did not consistently show stronger cross-species similarity in state frequency between human and mouse than pairs with low human-only baseline scores (**Supplementary Fig. 8**).

### LECIF score’s relationship to annotations based on sequence

We next analyzed the relationship between the LECIF score and various sequence-based annotations of conservation. We found that human regions overlapping sequence constrained elements called by GERP++^22^, SiPhy-omega, SiPhy-pi^23,51^, or PhastCons^20^ had a greater average LECIF score (0.19 to 0.22) than the genome-wide mean LECIF score (0.14) (**Fig. 5a**). We also compared the LECIF score to five sequence constraint scores and additionally the percent identity between human and mouse. For these comparisons, we slid a 50-bp window across the human genome and computed the mean of each score (LECIF or sequence constraint) in each window to control for resolution differences (**Fig. 5c, Supplementary Fig. 11**; **Methods**). The LECIF score was moderately correlated with sequence constraint scores, with PCCs ranging from 0.18 to 0.25 for windows with all bases annotated by the scores being compared. We note that this moderate correlation may partly reflect the incompleteness of functional genomics data or differences of the resolution of the data types we were not able to control for, in addition to conservation at the functional genomics level that is not represented in sequence constraint.

**Figure 5.**
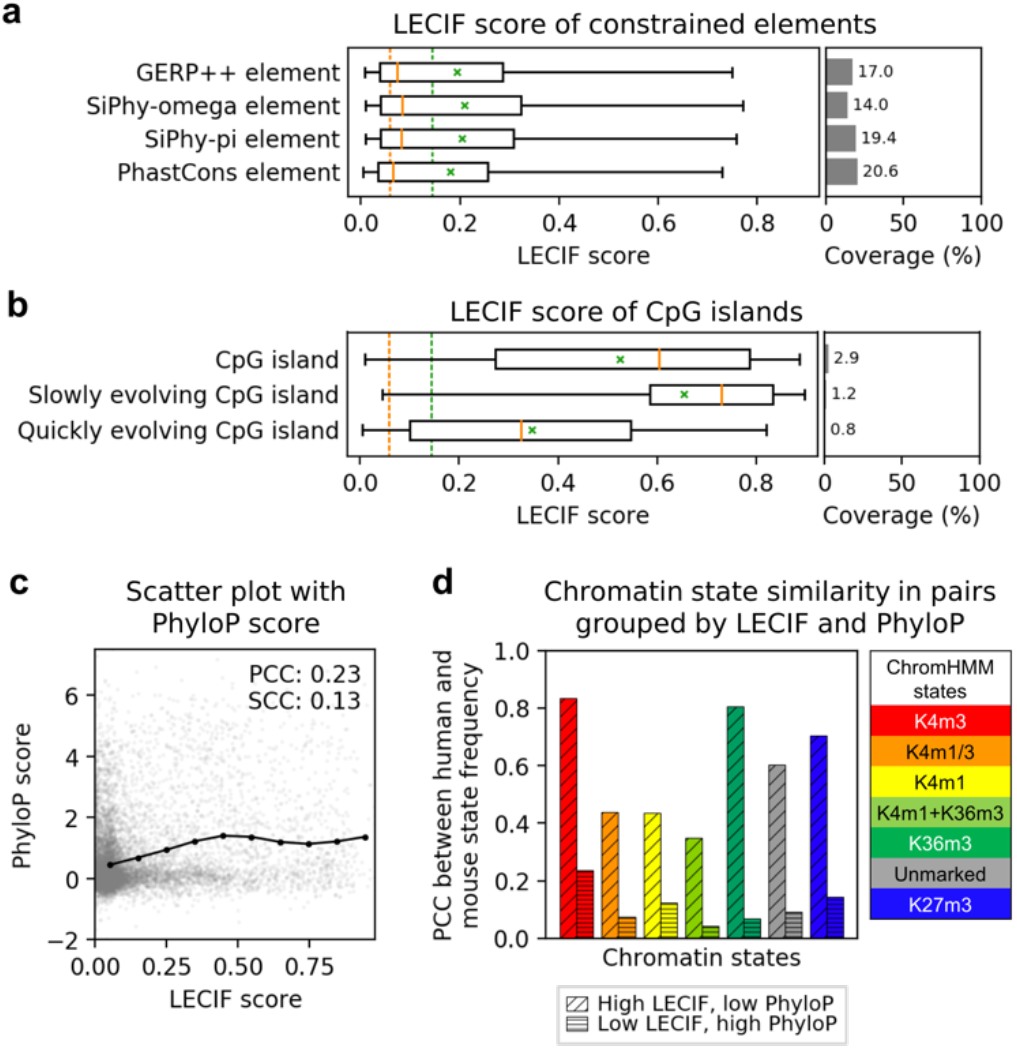
Relationship of LECIF score to sequence constraint annotations. **a.** Distribution of LECIF score within four different annotations of sequence constrained elements in human^20,22,23,51^. Each distribution is represented by a boxplot with median (orange solid line), mean (green ‘x’), 25^th^ and 75^th^ percentiles (box), and 5^th^ and 95^th^ percentiles (whisker). Dashed vertical lines in orange and green across the entire panel denote the genome-wide median and mean LECIF scores, respectively. Right sub-panel shows coverage of each annotation across all human regions that align to the mouse genome. **b.** Similar plot to **a**, except showing LECIF score of human regions overlapping any CpG island as well as subsets of slowly and quickly evolving CpG islands as previously defined based on primate sequence evolution^53^. **c.** Scatter plot showing with gray dots the LECIF score (x-axis) and PhyloP score^21^ (y-axis) for human regions. The PhyloP score was the human PhyloP defined based on a 100-way vertebrate alignment. The scatter plot is displaying 100,000 randomly sampled human regions that align to mouse and have all bases annotated by both scores. PCC and SCC, computed from all applicable human regions before downsampling, are shown in the top right corner. In black circles, the mean PhyloP score of all applicable human regions binned by the LECIF score with ten equal-width bins shown. These circles are connected with lines based on piecewise linear interpolation. **d.** ChromHMM chromatin state^24,44^ frequency correlation between human and mouse in pairs where the LECIF score is high and the PhyloP score is low or vice versa. The PhyloP score is the same as in **c**. The chromatin states are the same as in **Fig. 4b-d**. Bars with a diagonal hatch pattern show PCC computed from pairs with high LECIF score (>90^th^ percentile) and low PhyloP score (<10^th^ percentile) in all bases within 500 bp of the human region. Bars with a horizontal hatch pattern show PCC computed from pairs with low LECIF score (<10^th^ percentile) and high mean PhyloP score (>90^th^ percentile) in the human region. Bars for each state are colored according to the legend on the right. Similar plots with different percentile cutoffs and also including PCC for pairs that scored above the high or below the low thresholds for both scores are shown in **Supplementary Fig. 12**.

To provide evidence that high LECIF scores observed in regions with low sequence constraint scores are unlikely LECIF’s false positives, we analyzed human and mouse chromatin state annotations in regions where the two scores strongly disagreed. Specifically, for pairs of regions where the LECIF score was high and PhyloP score^21^ was low in all bases within 500 bp of the human region, we computed the correlation of chromatin state frequencies as described in the previous section (**Fig. 5d, Supplementary Fig. 12**). We found that such pairs had strong cross-species similarity for all and high PhyloP score had weaker cross-species similarity of frequency in all states. This suggests that the LECIF score can capture conservation at the functional genomics level even in regions that lack sequence constraint, potentially detecting signatures of conservation not captured by sequence constraint annotations.

We next investigated whether we could identify patterns within a multi-species sequence alignment that correspond to the differences between the LECIF score and constraint scores. To do this, we leveraged the ConsHMM^52^ 100-conservation-state annotation of the human genome based on a 100-way vertebrate sequence alignment (**Supplementary Fig. 13**). The conservation state with the highest average LECIF score was a state showing moderate probability of alignment and matching through many vertebrates, and previously shown to have the strongest enrichment for promoter and CpG islands. In contrast, this state had only the 12^th^ highest average PhyloP score. This suggests that some of the disagreement between the LECIF score and constraint scores could correspond to constraint scores not capturing signatures of conservation that are actually present in the multi-species sequence alignment.

We also analyzed the LECIF score in the context of annotations of human CpG islands previously grouped by their distinct regimes during primate sequence evolution^53^ (**Fig. 5b**). CpG islands in general scored high with a mean LECIF score of 0.53, and the score positively correlated with the likelihood of a CpG island being classified as slowly evolving as opposed to quickly evolving (**Supplementary Fig. 14;** PCC: 0.50). Slowly evolving CpG islands characterized by low rate of C-to-T deamination had higher LECIF scores with a mean of 0.65. In contrast, quickly evolving CpG islands had lower LECIF scores with a mean of 0.35.

### High-scoring regions are enriched for phenotype-associated variation and partitioned heritability of traits

We next investigated the relationship between the LECIF score and phenotype associated genetic variation (**Fig. 6a**). We observed that regulatory disease variants from Human Gene Mutation Database (HGMD)^54^ enriched for regions with high LECIF score. In contrast, we saw small depletions for common variants^55^ in those high-scoring regions. We saw that high-scoring regions also exhibited enrichment of Genome-wide Association Studies (GWAS) Catalog^56^ variants and expression quantitative trait loci (eQTLs) from GTEx^57^.

**Figure 6.**
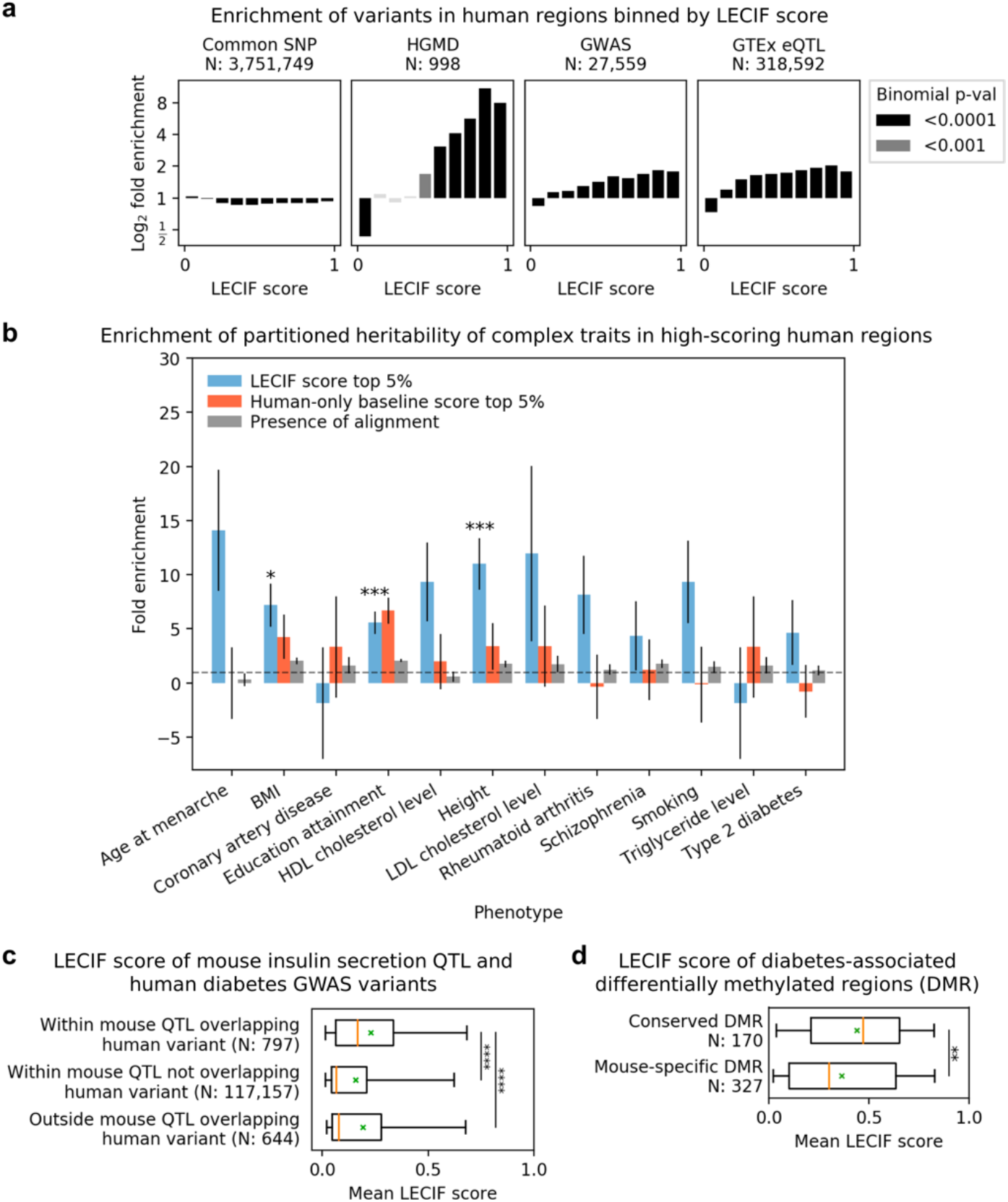
Relationship of LECIF score to genetic and epigenetic variation associated with phenotypes. **a.** Shown from left to right are plots of log_2_ fold enrichment for variants based on four different sets, (i) common SNPs^55^, (ii) HGMD regulatory variants^54^, (iii) GWAS catalog SNPs^56^, and (iv) GTEx cis-eQTLs^57^ across tissues, within human regions binned by the LECIF score with ten equal-width bins. Analysis was restricted to human regions that align to mouse, and a uniform background within these regions was used. Displayed above each subplot is the number of regions overlapping the variants from the corresponding set included in the analysis. Black and dark grey bars denote log_2_ fold enrichments that resulted in p-values below 0.0001 and 0.001, respectively, based on one-sided bionomial tests. **b.** Fold enrichments for partitioned heritability of 12 phenotypes^7^ in human regions with high LECIF score. Enrichments are shown for human regions with high human-mouse LECIF score (>95^th^ percentile) (blue) and additionally for comparison regions with high human-only baseline score (>95^th^ percentile) (orange) and human regions that align to mouse (gray). Heritability partitioning^7^ for the LECIF score was applied in the context of a baseline set of annotations^58^, which included sequence constraint annotations and was extended to include additional annotations generated based on the human-only baseline score and sequence alignment (**Methods**). Error bars denote jackknife standard error. Horizontal dashed line denote no enrichment (fold enrichment of 1). * and *** denote Bonferroni-corrected p-values for the LECIF score annotation’s enrichment below 0.05 and 0.001, respectively. **c.** Distribution of mean LECIF score of non-overlapping 1-kb human genomic windows identified as lying within a mapped mouse insulin scretion QTL or containing a human diabetes GWAS variant or both^59^. The top group on the plot, ‘Within mouse QTL overlapping human variant’, refers to windows that lie within the mouse QTL mapped to human and overlap the human diabetes GWAS variant. ‘Within mouse QTL not overlapping human variant’ refers to windows within the mouse QTL that do not overlap any human diabetes GWAS variant. ‘Outside mouse QTL overlapping human variant’ refers to windows randomly sampled from the human genome that lie outside the mouse QTL and overlap the human diabetes GWAS variant. Any window with less than half of its bases annotated with the LECIF score was excluded from this analysis. Displayed after each label is the number of qualified windows corresponding to that label. Each distribution is represented by a boxplot with median (orange solid line), mean (green ‘x’), 25^th^ and 75^th^ percentiles (box), and 5^th^ and 95^th^ percentiles (whisker). **** denotes p-value below 0.0001 based on a Mann-Whitney U test. Similar plots generated using different window sizes are shown in **Supplementary Fig. 15**. **d.** Distribution of mean LECIF score in conserved differentially methylated regions (DMRs) and mouse-specific DMRs with respect to a diabetic phenotype^60^. ‘Conserved DMR’ refers to regions with significant differential methylation (p-value<0.05) in both human and mouse and the same directionality with respect to the phenotype. ‘Mouse-specific DMR’ refers to regions with significant differential methylation in mouse, but either lacking significant differential methylation in human or showing inconsistent direction of methylation change between human and mouse. The study in which the DMRs were reported did not provide human-specific DMRs because it first identified mouse DMRs and then tested those in human and not vice versa. Displayed below each label is the number of DMRs corresponding to that label. Boxplots are formatted as in **c.**\*\* denotes p-value below 0.01 based on a Mann-Whintey U test.

We also conducted a heritability partitioning analysis with the LECIF score for 12 complex traits^7^. Specifically, we applied heritability partitioning with an annotation of bases with a LECIF score in the top 5% in the context of a baseline set of annotations^58^, which we extended to also include annotations of human regions that align to mouse and top 5% regions based on the human-only baseline score. We note that the baseline annotation set includes multiple sequence constraint annotations. Even in the context of this extended set of baseline annotations, we observed that the LECIF annotation resulted in enrichments of heritability with statistical significance for several traits (**Fig. 6b**). Furthermore, we observed overall stronger enrichments for the LECIF annotation than the human-only baseline annotation and the annotation of human regions that align to mouse. These results suggest that the LECIF score provides complementary information to sequence constraint annotations in elucidating the link between variants and phenotype.

### LECIF score highlights regions within mouse quantitative trait loci relevant to human disease

To demonstrate information captured by LECIF for translation of biological findings between mouse and human in a phenotypic context, we analyzed mouse insulin secretion quantitative trait loci (QTL) and human diabetes GWAS variants^59^. Previously, it was shown that human regions syntenic to the mouse insulin secretion QTL were enriched for the human diabetes GWAS variants. However, mouse QTL in general can span several megabases, making it difficult to identify likely causal variants within the loci for the trait of interest^18^. We thus asked whether the LECIF score could provide information in locating regions within the mouse insulin secretion QTL that correspond to human diabetes GWAS variants.

We observed that genomic windows within the mouse insulin secretion QTL that overlap the human GWAS variants had a statistically higher distribution of mean LECIF scores than windows within the mouse QTL not overlapping the variants or windows outside the mouse QTL overlapping the variants (Mann-Whitney U test *P*<0.0001; **Fig. 6c**, **Supplementary Fig. 15b,c**). Additionally, we saw that the human diabetes GWAS variants that lie within the mouse QTL had a higher distribution of mean LECIF scores than human GWAS variants outside the mouse QTL in addition to bases within the mouse QTL that are not the human GWAS variants (Mann-Whitney U test *P*<0.0001; **Supplementary Fig. 15a**). These results indicate LECIF’s potential value in finding genomic variants with phenotypic association in both species.

### LECIF score highlights regions with conserved differential methylation patterns linked to phenotype

We also evaluated the ability of the LECIF score to prioritize epigenetic features conserved between human and mouse in a disease relative context. Specifically, we considered data from an epigenetic study that examined differential methylation in diabetic phenotypes in both human and mouse^60^, which was independent of the data used to train LECIF. The study identified differentially methylated regions (DMRs) in high-fat-fed and low-fat-fed mouse adipocytes. The homologous human regions of the identified mouse DMRs were then tested to determine if they were differentially methylated as well between lean and obese patients, identifying regions with conserved differential methylation patterns associated with obesity in both human and mouse. The LECIF score was significantly higher in conserved DMRs in comparison to mouse-specific DMRs (Mann-Whitney U test *P*<0.01; **Fig. 6d**). This supports the potential value of the LECIF score for prioritizing among all loci with epigenetic associations with phenotype in one species the specific loci with assocations that are more likely to be shared in the other species.

## Discussion

We proposed LECIF, a method that scores evidence for conservation between human and mouse based on a compendium of functional genomic annotations from each species. To do so, LECIF trains neural networks to differentiate aligning pairs of regions from mismatched pairs of the same set of regions based on their functional genomic annotations without using sequence information as features. The functional genomic annotations include maps of open chromatin, transcription factor binding, gene expression signals, and chromatin state annotations. The resulting score captures evidence of conservation at the functional genomics level that is based on a diverse set of annotations and thus not specific to one class of DNA elements.

We applied LECIF with more than 10,000 functional genomic annotations from human and mouse to learn the human-mouse LECIF score. The LECIF score had greater predictive power than several baseline scores at discriminating pairs of human and mouse regions that align to each other from mismatched pairs of aligning regions. Using H3K27ac samples matched by their tissue of origin and separately using chromatin state annotations learned jointly between human and mouse, we showed that the LECIF score reflects the relationships between biologically similar human and mouse functional genomic annotations. LECIF was able to do this without any explicit information provided about the relationship between different features within or across species. Furthermore, LECIF was able to do so even in regions where sequence constraint was low, supporting that the LECIF score provides complementary information to sequence constraint annotations. Additionally, regions with high LECIF score were enriched for phenotype-associated variants, even when conditioning on sequence constraint annotations. Using matched DNA methylation samples between human and mouse and separately using matched GWAS and QTL data sets, both in the context of a diabetes trait, we showed that the LECIF score has preference for regions with shared associations with the trait. These results support the potential value of the LECIF score in various applications in the context of model organism research, such as highlighting genomic or epigenetic variation in mouse relevant to human traits or prioritizing human genomic variants to test in transgenic mice.

While we expect LECIF to be useful, we do note a few limitations. LECIF only scores evidence of conservation at the functional genomics level. Thus there could be regions that are conserved at the functional genomics level, but have a low LECIF score, since the evidence was not present in the data currently available to LECIF. Fortunately, the interpretation of high LECIF scores is less ambiguous. We also note that the LECIF score’s resolution is limited by the resolution of the input functional genomic annotations and thus does not have the base resolution that sequence-based conservation annotations can have.

Additionally, we note that currently the LECIF score is only available for pairs of regions that align to each other. While in principle LECIF can be applied to score any pairs of regions, more false positive predictions are expected as a result, compared to our presented strategy of restricting to regions that align at the sequence level. Although we explored an alternative strategy that considered non-aligning regions in a neighborhood of each pair of aligning regions, this did not lead to improvements in our evaluations over considering only the aligning regions. However, future work could develop other strategies that lead to improvements.

While here we focused on human and mouse, as mouse is a widely used model organism for human and there is substantial data available for both, LECIF can be applied to any pair of species though the quality of the score will depend on the coverage of the data available for both species. As functional genomic data from a more diverse set of species, cell types, and assays continues to become available, the utility of LECIF will continue to grow for identifying regions conserved at the functional genomics level and transferring findings from model organism research to human biology.

## Material and Methods

### Human-mouse LECIF score and code availability

The human-mouse LECIF score and the LECIF code is available at https://github.com/ernstlab/LECIF.

### Pairwise sequence alignment

For the pairwise sequence alignment, we used the chained and netted alignment^19^ between the human genome (hg19) and the mouse genome (mm10), with human as the reference genome for the alignment. This alignment was obtained from the UCSC Genome Browser^46^ (http://hgdownload.cse.ucsc.edu/goldenpath/hg19/vsMm10/axtNet/).

### Functional genomics data used for input features

ChromHMM^44^ chromatin state annotations for human were from the 25-state model learned for 127 cell and tissue types based on imputed data from the Roadmap Epigenomics Project^3^ and for mouse from the 15-state model learned for 66 cell and tissue types from ENCODE^61^. Peak calls for DNase-seq and ChIP-seq experiments of transcription factors, histone modifications, and variants were from Roadmap Epigenomics^3^, ENCODE^2^, and Mouse ENCODE^24^. Peak calls for Cap Analysis Gene Expression (CAGE) experiments were from FANTOM5^45^. RNA-seq signal data were from ENCODE^2^ and Mouse ENCODE^24^. For ENCODE and Mouse ENCODE data, we used the uniformed processed version available from the ENCODE portal. Additional information including the specific source of each dataset used is listed in **Supplementary Table 1**.

### Defining pairs of human and mouse regions for training and prediction

To define pairs of human and mouse regions for training and prediction for LECIF, we first identified alignment blocks from the pairwise alignment. We defined alignment blocks as pairs of human and mouse genomic segments without any alignment gap, meaning the human and mouse genomic segments both had a nucleotide present at each base in the block. We then for each alignment block defined non-overlapping windows of 50 bp starting from the first base in the alignment block. Each 50-bp window defined a region. If the alignment block ended within the 50-bp window, we truncated the window to the end of the block to define the region. This resulted in some regions being shorter than 50 bp. To define negative examples, we randomly paired up human and mouse regions included in the positive examples. With this procedure, all human regions included in the negative examples aligned somewhere else in the mouse genome, and all mouse regions in the negative examples aligned somewhere else in the human genome.

### Defining subsets of pairs of regions for training and evaluation

All human and mouse chromosomes, except for Y and mitochondrial chromosomes, were used. X chromosomes were excluded from training, validation, and test, but included for prediction and downstream analyses. To generate predictions for all pairs of human and mouse regions that included a human region from an even chromosome or X chromosome, we trained LECIF on pairs of human and mouse regions such that both the human and mouse regions came from a subset of odd chromosomes for its respective species (**Supplementary Table 2**). To form a validation set, which we used for hyper-parameter tuning and early stopping during training, we used pairs of regions such that the human region came from a subset of odd chromosomes not used in training and likewise for mouse. To form a test set, which we used to generate ROC and PR curves, we used all pairs of regions such that both the human and mouse region were from an even chromosome. To generate predictions for all pairs that included a human region from an odd chromosome, we took an analogous approach as above (**Supplementary Table 2**). There was no overlap in genomic regions used for training, validation, and test. To assess the agreement between a model trained on odd chromosomes and a model trained on even chromosomes, we used pairs of regions that were from a subset of chromosomes not used in training or validation of either model (**Supplementary Table 2**).

### Feature representations

For each pair of human and mouse regions, we generated two feature vectors. The two vectors were based on annotations overlapping the first base of the human and mouse regions, respectively, which were at most 50 bp. For computational considerations, we only used the first base of each region to provide the LECIF score for all bases in the region. However, we confirmed that if we had used the previously trained LECIF model to compute a score based on the annotations overlapping each base we would have learned a very similar score. Specifically, we computed the Pearson correlation coefficient (PCC) between a score defined at base resolution for 1 million randomly sampled pairs of human and mouse bases that align to each other and the LECIF score, which was defined at every 50 bp within each alignment block, for the same set of 1 million pairs.

Each peak call corresponded to one binary feature. If a base overlapped a peak call for an experiment, the corresponding value in the feature vector was encoded as a 1, otherwise it was encoded as a 0. Chromatin state annotations were one hot-encoded such that there was a separate binary feature representing the presence of each chromatin state in each cell or tissue type. Each RNA-seq experiment corresponded to one continuous feature. For human RNA-seq experiments, to also have the features in the range 0 to 1, we first computed the maximum and minimum signal value at any base in any of the human RNA-seq experiments. We then normalized values by subtracting the minimum signal value and dividing by the difference between the maximum and minimum signal values. We separately did the same normalization for mouse RNA-seq experiments.

### LECIF Classifier

The classifier that LECIF uses is an ensemble of neural networks where each neural network had a pseudo-Siamese architecture^62^ (**Supplementary Fig. 16**). A Siamese neural network consists of two identical sub-networks followed by a final sub-network that combines the output from the two sub-networks to generate a final prediction^63^. A pseudo-Siamese network is similar except it uses two distinct sub-networks instead of identical sub-networks. In LECIF, the two sub-networks corresponded to human and mouse. Human and mouse feature vectors were given to the human and mouse sub-networks, respectively, as input. We also evaluated using a fully-connected neural network, but found that it led to highly similar predictions (PCC: 0.95), but took longer to train.

Hyper-parameters of a neural network consisted of number of layers in each sub-network and the final sub-network, number of neurons in each layer, batch size, learning rate, and dropout rate. To set the values of the hyper-parameters, we conducted a random search, where we generated 100 neural networks, each with different randomly selected combinations of hyper-parameters (**Supplementary Table 3**). Each neural network was trained on the same set of randomly selected 1 million positive and 1 million negative training examples. We applied 50 times more weight to our negative examples than positive examples during training so that a high LECIF score corresponds to strong evidence of conservation. We identified the best-performing combination of hyper-parameters based on maximizing the AUROC on the validation examples.

With the best-performing combination of hyper-parameters, we then trained a new set of 100 neural networks each provided with different subsets of 1 million positive and 1 million negative training examples randomly selected from a pool of all training examples (>2.2 million positive and >2.2 million negative). We applied the same increased weighting of negative examples as above. The final prediction of the ensemble was the average of the predictions from the 100 trained neural networks.

For both hyper-parameter search and training, we stopped training if there were no improvements in AUROC evaluated on the validation examples over three epochs. We saved the classifier from the epoch with the highest AUROC on the validation examples. The maximum number of epochs we allowed during training was 100 and the maximum training time we allowed was 24 hours.

We also generated a version of the LECIF classifier, LECIF-GB, which was trained in the same way as LECIF except the negative examples were pairs of human and mouse regions that were both randomly selected from anywhere in their respective genomes as opposed to being constrained to aligning regions.

We used PyTorch (version 0.3.0.post4)^64^ for implementation of the neural networks.

### Random forest baseline

We trained, applied, and evaluated random forest using the same procedure as explained above, except we used a decision tree in place of a neural network. We also did hyper-parameter selection as explained above, but for a set of hyper-parameters unique to decision trees (**Supplementary Table 3**). We used Scikit-learn (version 0.19.1)^65^ for implementation.

### Canonical correlation analysis baseline

We trained an ensemble of canonical correlation analysis (CCA) mappings using the same procedure as above, except using a CCA mapping in place of a neural network and positive examples only. We applied and evaluated the ensemble using the same procedure as explained above. We also did hyper-parameter selection as explained above, but for a set of hyper-parameters unique to CCA mapping (**Supplementary Table 3**) and through a grid search instead of random search. We used Pyrcca^66^ for implementation.

### Logistic regression baseline

We trained, applied, and evaluated an ensemble of logistic regression classifiers using the same procedure as above, except we used a logistic regression classifier in place of a neural network. We also did hyper-parameter selection as for the neural networks, but for a set of hyper-parameters unique to logistic regression models (**Supplementary Table 3**) and through a grid search instead of random search. We used Scikit-learn (version 0.19.1)^65^ for implementation.

### Human-only baseline

We trained, applied, and evaluated a human-only baseline, which used the same human features as LECIF, but did not use any mouse features. For training the human-only baseline, positive examples were human regions that align to the mouse genome and the negative examples were human region that do not align to the mouse genome. We otherwise used the same procedure for training, prediction, and evaluation as for LECIF except we used an ensemble of fully-connected neural networks. We also did hyper-parameter selection as for LECIF, but for a set of hyper-parameters of a fully-connected neural network (**Supplementary Table 3**). We used PyTorch (version 0.3.0.post4)^64^ for implementation.

### Computing area under the receiver operating characteristic and precision recall curves

To compute each classifier’s classification performance based on area under the receiver operating characteristic curve and precision-recall curve, we used Scikit-learn’s implementation^65^.

### Defining a region-neighborhood LECIF score

To generate a region-neighborhood LECIF score for each pair of a human region and its aligning mouse region, we computed LECIF scores for additional pairs that consisted of the same human region and distinct 50-bp mouse regions located within a neighborhood of *W* bases centered around the aligning mouse region **(Supplementary Fig. 3)**. The non-aligning mouse regions were defined by sliding a 50-bp window from the first base of the aligning mouse region in both the 5’ and 3’ directions. We then took the maximum over these LECIF scores to produce the region-neighborhood LECIF score. We varied *W* between 0 and 20kb. We note that *W* of 0 corresponds to the original LECIF score.

### H3K27ac activity similarity

To define the H3K27ac activity similarity between human and mouse based on known biology, we took all human and mouse H3K27ac experiments used for features and manually grouped them into the following 14 tissue type groups based on available annotations of the experiments: adipose, bone element, brain, embryo, heart, intestine, kidney, limb, liver, lung, lymph node, spleen, stomach, thymus. **Supplementary Table 1** specifies which experiment was assigned to which group, but we note that information about these groups were not used in learning the LECIF score. The 14 groups listed above were represented in at least one H3K27ac experiment in both species. For the analysis, we discarded experiments that did not belong to any of the tissue groups.

For each pair of human and mouse regions, we then defined vectors *h* and *m* of length 14 where *h*_*i*_ and *m*_*i*_ correspond to the fraction of experiments in the *i*-th group with peak calls that overlapped the human and mouse regions, respectively. Finally, for each pair of human and mouse regions, we computed the weighted Jaccard similarity coefficient^67^ between these two vectors. The weighted Jaccard similarity coefficient is defined as:

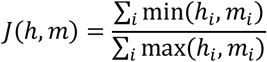

Any pair with an undefined similarity coefficient, due to the denominator summing up to zero was removed from the analysis.

### Chromatin state frequency correlation

To analyze cross-species agreement of chromatin state frequencies as a function of the LECIF score, we first grouped pairs of human and mouse regions based on their LECIF score. When binning based on either score, five or ten equal-width bins were used with varying numbers of pairs in each bin. We repeated the procedure when using the human only baseline score in place of the LECIF score. We also binned based on the percentile rank of scores, where either five or ten bins were used with nearly the same number of pairs in each bin.

To compute the chromatin state frequency correlation across a set of pairs of human and mouse regions defined as described above, we used a chromatin state model jointly learned from both human and mouse genome^24^. For each of the seven chromatin states, we defined vectors for human and mouse. An element of a vector for human corresponds to the fraction of epigenomes in which one of the human regions is annotated with the state, and similarly for the mouse vector and regions. We then computed the PCC between the two vectors for each chromatin state, resulting in 7 PCC values.

### Correlation between the LECIF score and sequence constraint scores

To compute the correlation between the LECIF score and sequence constraint scores, we slid a 50-bp genomic window in 10-bp increment across the human genome. For each window, we computed the mean of each score (LECIF or sequence constraint). For each sequence constraint score, we computed the PCC and SCC between the LECIF score and the sequence constraint score for windows with at least *n* bases annotated by the two scores, with *n* ranging from 1 to 50. The two scores were not required to be defined on the same set of bases within the 50-bp window.

### Heritability partitioning analysis

To perform the heritability partitioning analysis, we used the LD-score regression software ldsc (v1.0.0)^7^. We generated an annotation of all human regions that align to the mouse genome and have a LECIF score above the 95^th^ percentile. We used this annotation in the context of the baseline annotation set from Gazal et al.^58^ along with another annotation generated based on the human-only baseline score instead of the LECIF score and an annotation of human regions that align to the mouse genome. We also included 500-bp windows around each annotation to dampen the inflation of heritability in neighboring regions due to linkage disequilibrium, following the procedure in Ref. 7.

We applied ldsc to this extended set of annotations for the following 12 traits: age at menarche, body mass index (BMI), coronary artery disease, education attainment, HDL cholesterol level, height, LDL cholesterol level, rheumatoid arthritis, schizophrenia, smoking, triglyceride level, and type 2 diabetes. Below we list the source of all summary statistics used in this analysis:

- Age at menarche^68^: https://www.reprogen.org
- Body mass index, height^69^: http://www.broadinstitute.org/collaboration/giant/index.php/GIANT_consortium_data_files
- Coronary artery disease^70^: http://www.cardiogramplusc4d.org/data-downloads
- Education attainment^71^: https://www.thessgac.org/data
- HDL cholesterol level, LDL cholesterol level, triglyceride level^72^: http://csg.sph.umich.edu/willer/public/lipids2010
- Rheumatoid arthritis^73^: http://plaza.umin.ac.jp/yokada/datasource/software.htm
- Schizophrenia^74^, smoking^75^: www.med.unc.edu/pgc/downloads
- Type 2 diabetes^76^: http://www.diagram-consortium.org/downloads.html

### Genic, regulatory, and variant annotations

For TSS, gene body, intron, exon, coding exon, 5’ UTR, and 3’ UTR annotations, we used GENCODE annotations V31lift37 for human and VM23 for mouse. We downloaded these annotations along with classification of evolutionary dynamics of CpG islands^53^ and common SNPs (dbSNP v7)^55^ from the UCSC Table Browser^46^. The HGMD variants that we used were variants annotated as ‘regulatory mutations’ in the April 2012 public release of HGMD database^6,54^.

## Acknowledgements

We thank the ENCODE, Mouse ENCODE, Roadmap Epigeomics, and FANTOM consortia for generating the data and making it publically available. We thank members of the Ernst lab for useful discussions. We acknowledge funding from US National Institutes of Health (DP1DA044371, U01MH105578 to J.E.); US National Science Foundation (CAREER Award #1254200 to J.E.); Kure It cancer research (Kure-IT award to J.E.), and a Rose Hills Innovator Award (J.E.).

