## Supplementary figures for "Learning a genome-wide score of human-mouse conservation at the functional genomics level"


1. Effect of different weight ratios between positive and negative examples
2. Effect of ensembling and sampling training data on robustness
3. Overview of generating region-neighborhood LECIF score for pairs of human regions and extended mouse regions
4. Predictive power of region-neighborhood LECIF score for aligning pairs as a function of neighborhood size around each pair's mouse region
5. Distribution of mean LECIF score of mouse peak calls provided to LECIF
6. Distribution of mean LECIF score in different mouse chromatin states
7. Distribution of LECIF score of mouse GENCODE gene feature annotations
8. Cross-species similarity in chromatin states in pairs binned by LECIF score or human-only baseline score
9. Relative frequency of chromatin states in regions with low or high LECIF score
10. Scatter plot of the human-only baseline score and cross-species similarity in tissue-specific H3K27ac activity
11. Correlation between LECIF score and sequence constraint scores
12. Cross-species agreement in chromatin state frequency in pairs grouped based on LECIF score and PhyloP score
13. Relationship of LECIF score and PhyloP score in ConsHMM conservation states
14. Relationship of LECIF score and log-odds score for CpG island being classified as slowly evolving
15. Distribution of mean LECIF score of human genomic windows overlapping mouse insulin secretion QTL and human diabetes GWAS variant
16. A schematic of a pseudo-Siamese neural network


**Supplementary Tables**


1. Feature summary, type, source, and metadata
2. Data split for training and evaluation
3. Hyper-parameter values considered and chosen for all methods

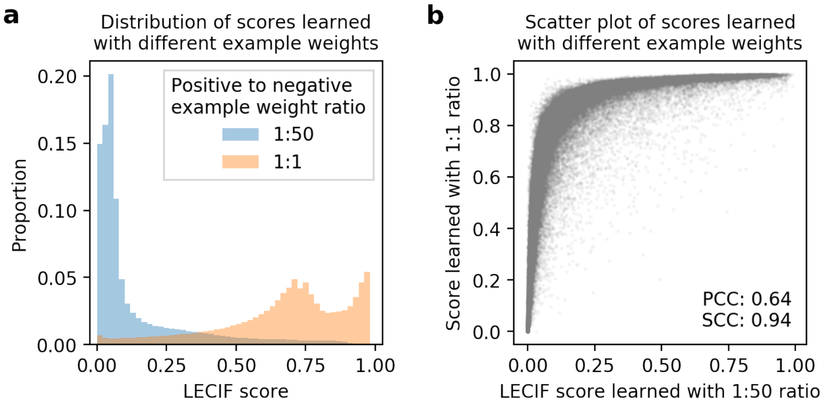


**Supplementary Figure 1. Effect of different weight ratios between positive and negative examples.**

A comparison of (i) the LECIF score, which was learned with negative examples weighted 50 times more than positive examples and (ii) an alternative version of the score learned with positive and negative examples weighted equally. To generate the alternative version, we repeated the hyper-parameter search and prediction procedures with the same dataset, but with equal weigthing scheme.

**a.** Distribution of the two scores. Blue bars correspond to the LECIF score. Orange bars correspond to the alternative version of the score learned with equal weights. Fifty equal-width bins were used for both scores to plot this histogram.

**b.** Scatter plot showing with a gray dot for each aligning pair of human and mouse regions the LECIF score (x-axis) and the alternative version of the score learned with equal weights (y-axis). Pearson correlation coefficient (PCC) and Spearman correlation coefficient (SCC) between the two scores are shown in the bottom right. One hundred thousand pairs of human and mouse regions were randomly selected to be included in the scatter plot.

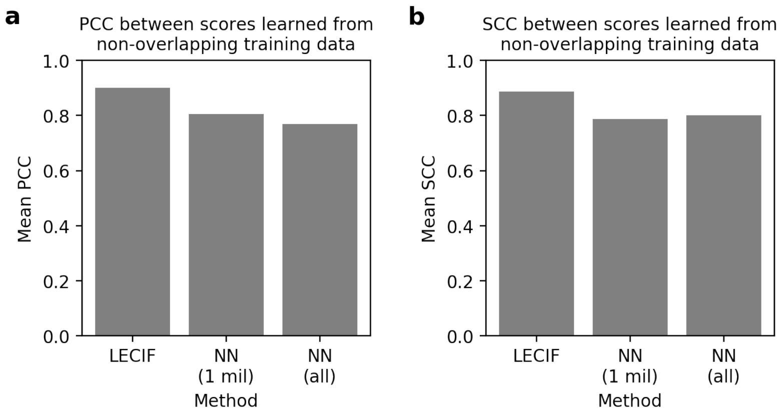


**Supplementary Figure 2. Effect of ensembling and sampling training data on robustness.** Analysis of the effect of the ensembling strategy of LECIF, which trains an ensemble of 100 neural networks, where each neural network (NN) is given 1 million positive and 1 million negative examples that are randomly sampled from all available training data, on the robustness of predictions. LECIF's robustness is compared the average robustness of individual NN in the ensemble and also the robustness of a single NN trained on all available training data (>2.2 million positive and >2.2 million negative examples). We measure the robustness by computing the **a.** PCC and **b.** SCC between scores generated by classifiers that were trained on non-overlapping set of chromosomes (**Methods**). 'NN (1 mil)' refers to the mean of 100 PCC or SCC computed for each of the 100 NN in the ensemble used in LECIF. 'NN (all)' refers to the PCC or SCC computed for a NN trained on all available training data. The scores we compare here were generated for pairs of human and mouse regions held out from training and validation (**Supplementary Table 2**). The same set of scores were used to compute both PCC and SCC. These results confirm that LECIF leads to more robust predictions than any individual NN in the ensemble or a NN trained on all available data.

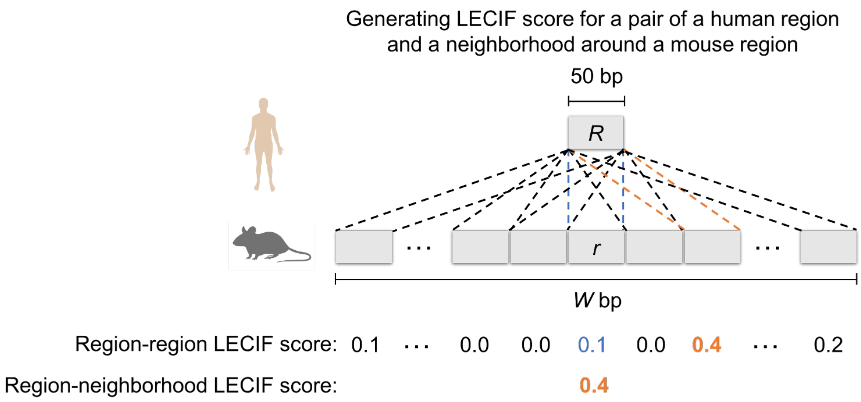


**Supplementary Figure 3. Overview of generating region-neighborhood LECIF score for pairs of human regions and extended mouse regions.**

Illustration of how LECIF is used to generate region-neighborhood LECIF score for a pair of a human region and a neighborhood of a mouse region. A given 50-bp human region *R* is compared to a set of multiple 50-bp mouse regions in a neighborhood of length *W* bp centered around a mouse region *r*. Each comparison (pair of dashed lines) results in a region-region LECIF score. For a pair of human region and a neighborhood in mouse, we define the region-neighborhood LECIF score as the maximum of all the region-region LECIF scores. In this example, the region-region LECIF score of the aligning human and mouse regions (blue; *R* and *r*) is 0.1. The maximum region-region LECIF score, 0.4, comes from the human region paired up with a mouse region near the aligning mouse region (orange). As a result, in this example, the region-neighborhood LECIF score is 0.4. We evaluated using the region-neighborhood LECIF score to predict aligning pairs, as an alternative to using the region-region LECIF score of the aligning human and mouse regions. Results of the evaluation are shown in **Supplementary Fig. 4**.

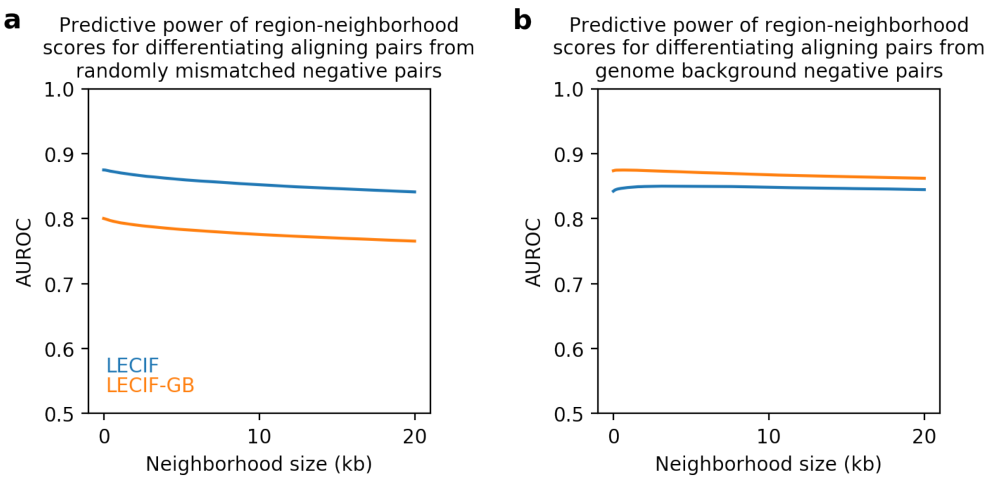


**Supplementary Figure 4. Predictive power of region-neighborhood LECIF score for aligning pairs as a function of neighborhood size around each pair's mouse region.**

We evaluate the predictive power of the region-neighborhood LECIF score of aligning human and mouse regions as a function of neighborhood size. We also evaluate using a LECIF-Genome Background (LECIF-GB) score in place of LECIF score in this analysis. LECIF-GB was trained with 'genome background' negative examples, which are pairs of human and mouse regions randomly selected from the entire human and mouse genomes (**Methods**). Shown for LECIF (blue) and LECIF-GB (orange) is the area under the ROC curve (AUROC) for differentiating positive examples from negative examples as a function of the size of the neighborhood centered around each pair's mouse region. Positive examples are pairs of human and mouse regions that align to each other. Negative examples are either **a.** randomly mismatched human and mouse regions that align somewhere in the other species (equivalent to the negative examples provided to LECIF) or **b.** genome background (equivalent to the negative examples provided to LECIF-GB). The neighborhood size varies from 0 to 20 kb with increments of 100 bp. Given a particular neighborhood size of *W*, the region-neighborhood score for each pair of human and mouse regions was the maximum region-region scores of any pair consisting of the human region and any mouse region within 0.5**W* bp from the aligning mouse region of the pair (**Methods**; **Supplementary Fig. 3**). This region-neighborhood LECIF score was then used to predict aligning pairs. We note that a neighborhood size of 0 gives region-region LECIF and LECIF-GB scores. For each comparison, the same set of 100,000 positive and 100,000 negative test examples, which were on chromosomes excluded from training and validation, were used to compute the AUROC. In this analysis, there was no advantage in using the region-neighborhood LECIF score, as defined, compared to using the region-region LECIF score and similarly for LECIF-GB.

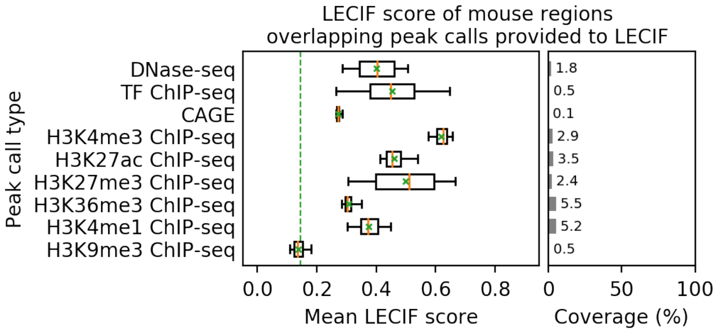


**Supplementary Figure 5. Distribution of mean LECIF score of mouse peak calls provided to LECIF.**

Similar to **Fig. 3a** except for mouse experiments instead of human**.** Shown for each type of functional genomic experiments listed is the distribution of mean LECIF score over experiments of that type in mouse. The mean LECIF score for an experiment is computed by averaging the LECIF score of mouse regions overlapping a peak call from the experiment. The set of experiments are the same as provided to LECIF as input features. Each distribution is represented by a boxplot with median (orange solid line), mean (green 'x'), 25$^{th}$ and 75$^{th}$ percentiles (box), and 5$^{th}$ and 95$^{th}$ percentiles (whisker). Green dashed vertical line across the entire panel denotes the genome-wide mean LECIF score. Right panel shows mean coverage of each type of peak call across all mouse regions that align to the human genome.

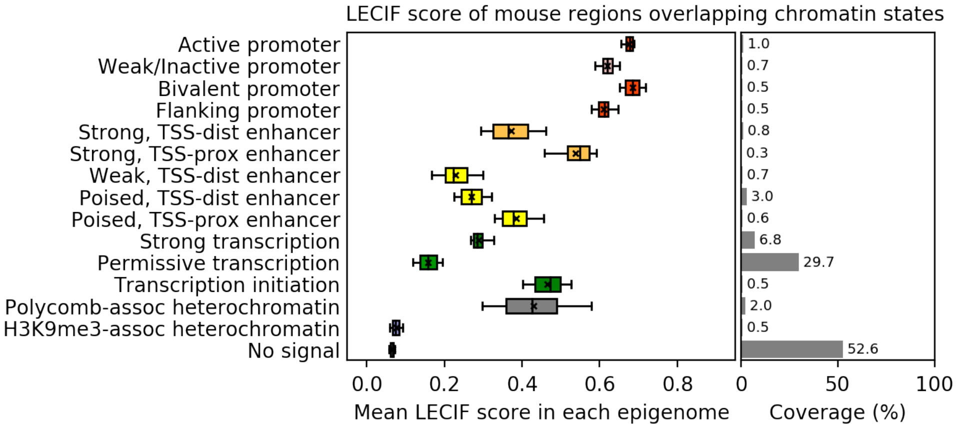


**Supplementary Figure 6. Distribution of mean LECIF score in different mouse chromatin states.** Similar to **Fig. 3b** except for mouse chromatin state annotations[1,2] instead of human**.** Shown for each chromatin state from a model learned in mouse is the distribution of mean LECIF score over different epigenomes. The mean LECIF score for a chromain state in an epigenome is computed by averaging the LECIF score of regions overlapping the chromatin state in the epigenome. Each distribution is represented by a boxplot with median (black vertical line), mean (black 'x'), 25$^{th}$ and 75$^{th}$ percentiles (box), and 5$^{th}$ and 95$^{th}$ percentiles (whisker). Right panel shows mean coverage of each state across all mouse regions that align to the human genome. State colors were assigned to match the state colors of the 25-state human ChromHMM model[3] shown in **Fig. 3b** based on state descriptions.

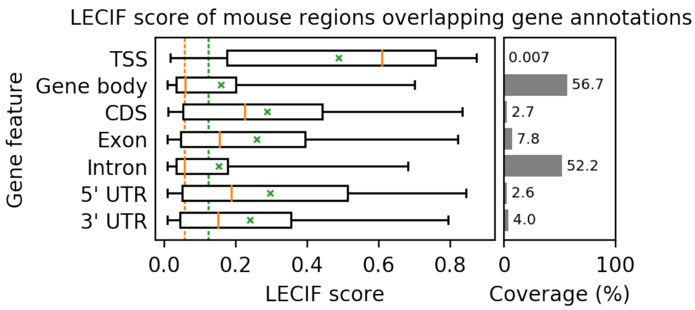


**Supplementary Figure 7. Distribution of LECIF score of mouse GENCODE gene feature annotations.**

Similar to **Fig. 3c** except for mouse regions overlapping mouse gene feature annotations instead of human. Distribution of LECIF score in mouse regions overlapping indicated GENCODE gene feature annotations. Each distribution is represented by a boxplot with median (orange solid line), mean (green ‘x’), 25$^{th}$ and 75$^{th}$ percentiles (box), and 5$^{th}$ and 95$^{th}$ percentiles (whisker). Orange and green dashed lines vertical lines across the entire panel denote the genome-wide median and mean LECIF scores, respectively. Right panel shows coverage of each annotation across all mouse regions that align to the human genome. TSS: transcription start site; CDS: coding sequence; UTR: untranslated region.

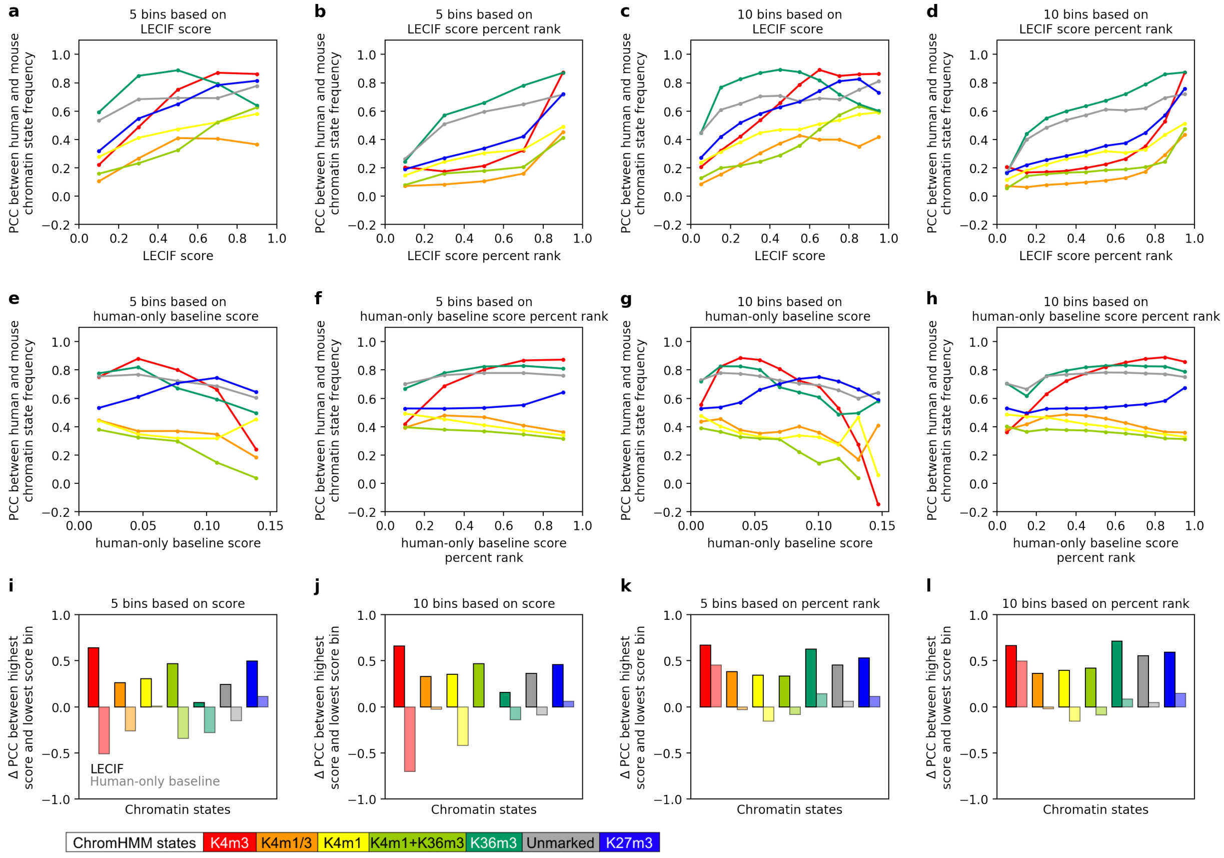


**Supplementary Figure 8. Cross-species similarity in chromatin states in pairs binned by LECIF score or human-only baseline score.**

Extended version of the analysis in **Fig 4b**. **a-d.** Cross-species agreement in chromatin state[2,4] frequency in pairs of aligning human and mouse regions binned by the LECIF score for a ChromHMM model learned jointly between human and mouse. Pairs are binned using **a.** 5 equal-width bins based on the LECIF score, **b.** 5 bins based on the percentile rank of the LECIF score, **c.** 10 equal-width bins based on the LECIF score, or **d.** 10 bins based on the percentile rank of the LECIF score. Binning based on the percentile rank results in similar number of pairs in each bin, whereas binning based on the score results in varying number of pairs in each bin. For each state and aligning region, we computed the frequency of the state across cell and tissue types for human and mouse separately. We then, for each state and bin, computed the PCC between the corresponding human and mouse frequencies for that state across all aligning pairs within the bin (**Methods**). The values are shown with colored circles according to the chromatin state legend on the bottom from Ref. 4. The circles for the same state are connected with lines based on piecewise linear interpolation. **d** is identical to **Fig. 4b**.

**e-h.** Similar to **a-d**, respectively, except using the human-only baseline score instead of the LECIF score.

**i-l.** Shown for each chromatin state (x-axis) is the difference in the chromatin state's PCC between pairs from the highest score and lowest score bin $\Delta$(PCC), based on either the LECIF score (bold-colored bars) or human-only baseline score (light-colored bars). Each panel corresponds to the two sub-panels above it in the same column. The $\Delta$PCC values are shown with colored bars according to the chromatin state legend on the bottom from Ref. 4 and the score used for binning the pairs (bold for LECIF score, light for human-only baseline score).

This figure illustrates that pairs of human and mouse regions with high LECIF score show stronger cross-species agreement in chromatin state frequency than pairs with low LECIF score. It also highlights that pairs with high human-only baseline score do not consistently show stronger cross-species agreement than pairs with low human-only baseline score.

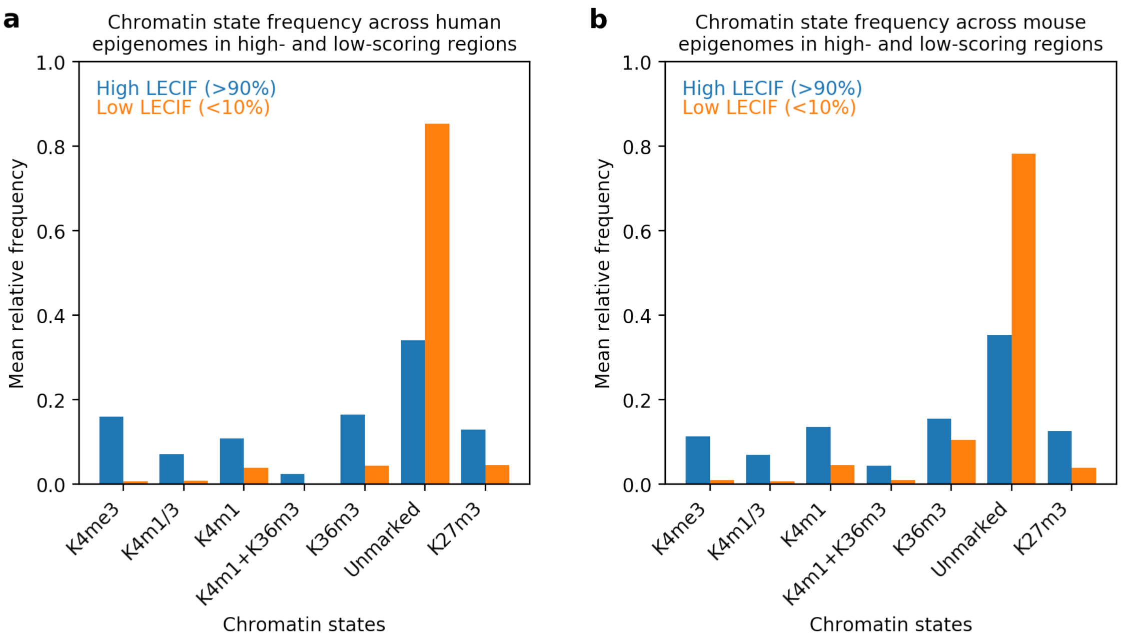


**Supplementary Figure 9. Relative frequency of chromatin states in regions with low or high LECIF score**.

Comparing relative frequency of chromatin states[2,4] for a seven state ChromHMM model learned jointly between human and mouse in high LECIF score (>90$^{th}$ percentile; blue) and low LECIF score (<10$^{th}$ percentile; orange) regions. The comparison is shown both for **a.** human and **b.** mouse regions. The chromatin states are the same as in **Fig. 4** and **Supplementary Fig. 8**. For a species, the mean relative frequency of a chromatin state in a set of regions satisfying the LECIF score threshold was computed by averaging over epigenomes the fraction of those regions overlapping the chromatin state in each epigenome. These figures illustrate that regions with low LECIF score are more likely to be annotated with the ‘Unmarked’ chromatin state in both human and mouse than regions with high LECIF score.

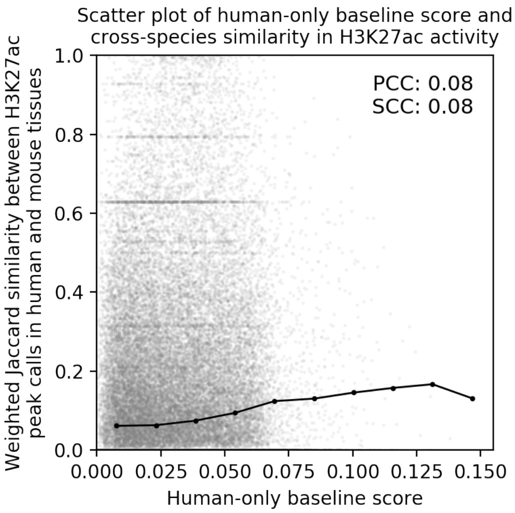


**Supplementary Figure 10**. **Scatter plot of human-only baseline score and cross-species similarity in tissue-specific H3K27ac activity.**

Similar scatter plot to **Fig. 4a** except for the human-only baseline score instead of the LECIF score. Scatter plot shows with a gray dot for each aligning pair of human and mouse regions the human-only baseline score (x-axis) and cross-species similarity of matched tissue-specific H3K27ac activity (y-axis). The H3K27ac activity for a region in a tissue and species is quantified as the fraction of experiments in the tissue type of the species with peak calls overlapping the region. The cross-species similarity of the tissue-specific H3K27ac activity is quantified as the weighted Jaccard similarity coefficient over 14 matched tissue types (**Methods**). PCC and SCC computed from all aligning pairs are shown in the top right. In black circles the mean similarity coefficient of pairs binned by the LECIF score with ten equal-width bins spanning from the minimum to maximum of the human-only baseline score is shown. These circles are connected with lines determined based on piecewise linear interpolation. One hundred thousand random aligning pairs were sampled to plot the scatter plot. This analysis shows that the human-only baseline score exhibits weaker agreement with cross-species similarity in tissue-specific H3K27ac activity compared to the LECIF score (PCC: 0.08 vs 0.45 and SCC: 0.08 vs 0.42).

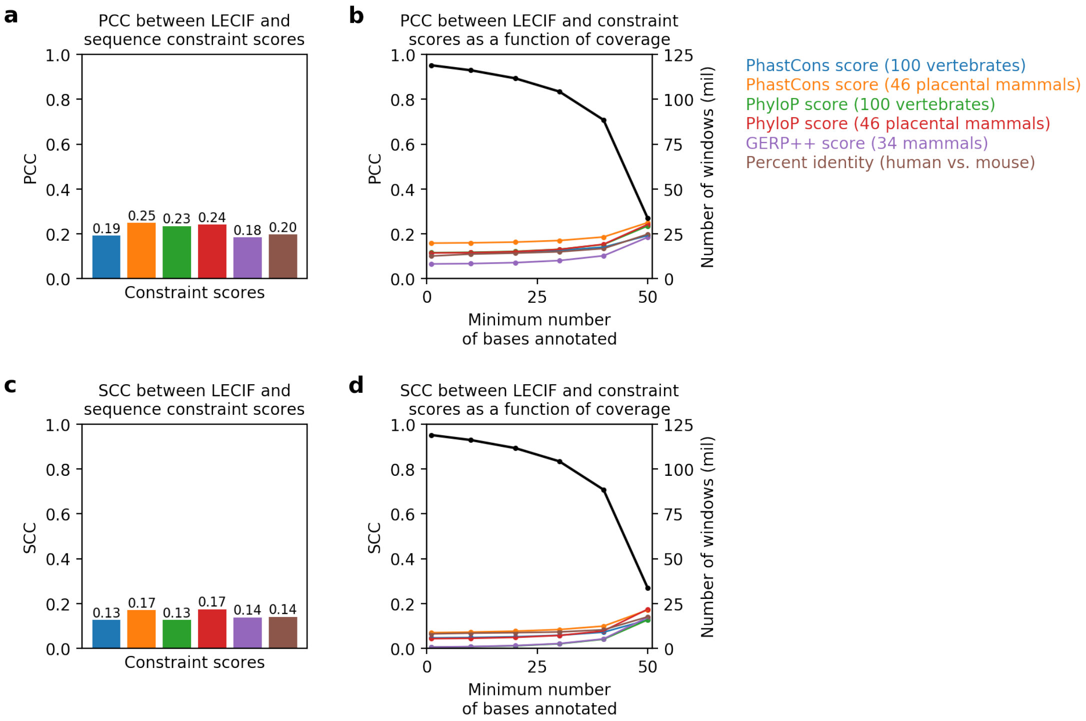


**Supplementary Figure 11. Correlation between LECIF score and sequence constraint scores.**
**a.** Shown for a set of sequence constraint scores[5–7] is the PCC computed between the LECIF score and a constraint score. For a given constraint score, to compute the PCC, we first slid a non-overlapping 50-bp genomic window across the human genome and selected windows with all 50 bases annotated by both the LECIF score and the given constraint score. We then computed the mean LECIF score and mean constraint score for each selected window. The PCC for a constraint score is the PCC between those two sets of values. Each resulting PCC is shown with a bar colored according to the legend on the right. Percent identity is defined as the number of matching base-pairs (e.g. G in human and G is mouse) within a given window divided by 50.
**b.** PCC between the LECIF score and constraint scores as a function of the minimum number of bases required to be annotated in the genomic windows. Also shown is the number of windows selected to compute the PCC. The PCC for a constraint score is computed as described in **a,** except windows with at least *n* bases annotated by the LECIF score and the constraint score of interest are selected, where *n* varies from 1 to 50. The two scores being compared need not annotate the same set of bases in each window. The PCC are shown with colored circles according to the y-axis on the left and legend in the top right. The circles for the same constraint score are connected with lines based on piecewise linear interpolation. The rightmost values where the minimum number of bases annotated equals 50 correspond to the PCC shown in **a**. Black circles show the number of windows in millions that had at least *n* bases (x-axis) annotated by the LECIF score and constraint scores according to the y-axis on the right. These circles are connected with lines based on piecewise linear interpolation. All six comparisons of the LECIF score to constraint scores had the same number of selected genomic windows.
**c-d.** Similar to **a-b**, respectively, except for SCC instead of PCC.
These results show that the LECIF score is moderately correlated with sequence constraint scores, and that the correlations are weaker as we allow fewer bases to be annotated by the scores within each genomic window.

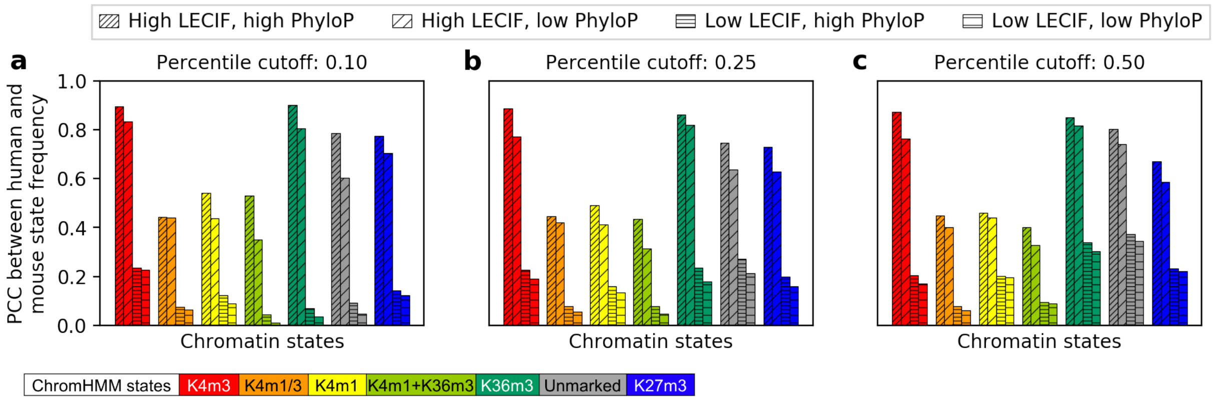


**Supplementary Figure 12. Cross-species agreement in chromatin state frequency in pairs grouped based on LECIF score and PhyloP score.**

**a.** ChromHMM chromatin state[4,8] frequency correlation between human and mouse in pairs of aligning human and mouse regions grouped based on whether their LECIF score and human PhyloP score[5] (defined based on a 100-way vertebrate alignment) were high (>90$^{th}$ percentile) or low (<10$^{th}$ percentile). The chromatin states are the same as in **Supplementary Fig. 8**. Separate bars are shown for each combination of high or low score of LECIF or PhyloP as indicated based on the legend at top. For the low PhyloP case, we required that there be a low (<10$^{th}$ percentile) score at all annotated bases within 500 bp to ensure the low score was not driven by the higher resolution at which sequence conservation is defined. The frequency correlation for each state and a set of aligning pairs is quantified as the PCC between the human and mouse frequencies for that state across the pairs, as done in **Supplementary Fig. 8** (**Methods**). Any region that did not have a PhyloP score for all bases was discarded from this analysis. Bars for each state are colored according to the bottom legend, as previously defined in Ref. 4.

**b.** Similar to **a** except using a percentile cutoff of 0.25 instead of 0.05. Scores above the 75$^{th}$ percentile are considered high, and scores below the 25$^{th}$ percentile are considered low.

**c.** Similar to **a** except using a percentile cutoff of 0.50 instead of 0.05. Scores above the median are considered high, and scores below the median are considered low.

These results demonstrate that pairs with high LECIF score exhibit strong cross-species agreement in chromatin state frequency even when there is a low PhyloP score in the region. In contrast, pairs with a high PhyloP score and a low LECIF score did not exhibit strong correlations.

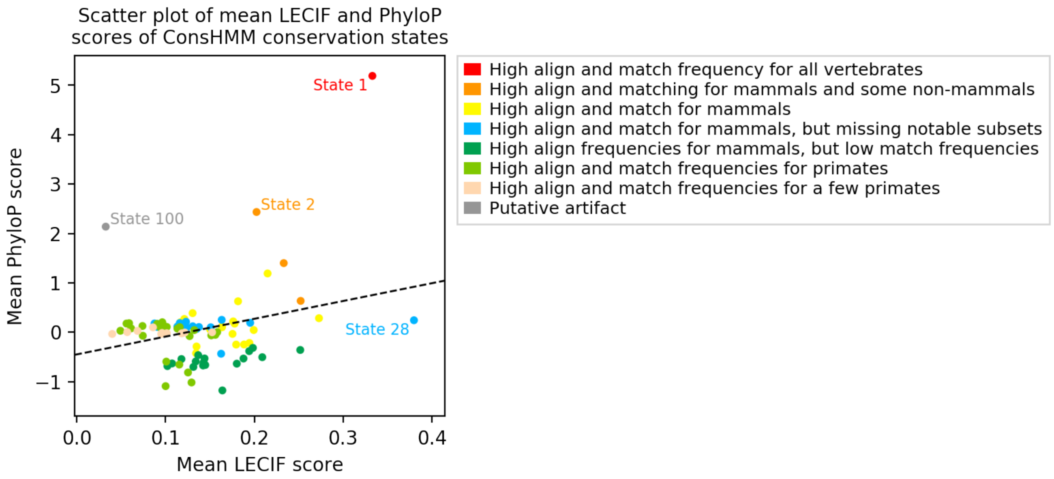


**Supplementary Figure 13. Relationship of LECIF score and PhyloP score in ConsHMM conservation states.**

We use a ConsHMM 100-conservation-state annotation of the human genome based on a 100-way vertebrate sequence alignment[9] to understand the relationship between the LECIF score and sequence constraint scores. The scatter plot shows with a dot for each ConsHMM conservation state the mean LECIF score (x-axis) and mean human PhyloP score[5] (y-axis; defined based on a 100-way vertebrate alignment). For each conservation state, the mean LECIF or PhyloP score is computed by averaging the score of bases overlapping the conservation state. Each dot is colored according to the eight major groups of conservation states listed in the legend on the right, as previously defined in Ref. 9. Dashed line is a linear regression fit applied to the 100 data points. We label four noteworthy conservation states. State 28 (blue), which is the promoter enriched state, has the highest mean LECIF score and the 12th highest mean PhyloP score. State 1 (red), which is the most enriched state for exons, has the 2nd highest mean LECIF score and the highest mean PhyloP score. State 2 (orange), which is the state most enriched for enhancer chromatin states, has the 8th highest mean LECIF score and the 2nd highest mean PhyloP score. State 100 (grey), which is characterized by pseudogenes and putative artifacts in the multi-species sequence alignment[9], has the lowest mean LECIF score, while having the 3rd highest mean PhyloP score.

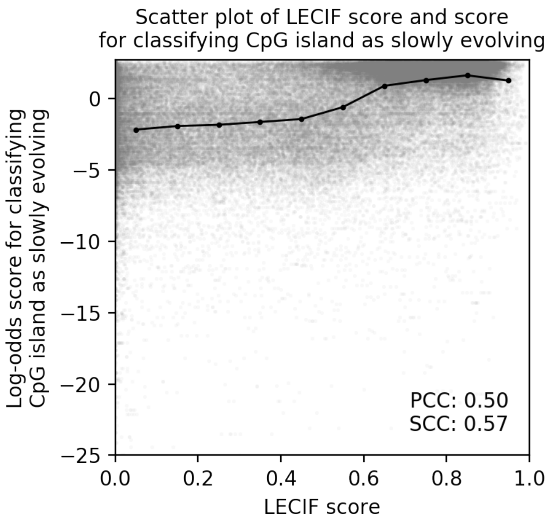


**Supplementary Figure 14. Relationship of LECIF score and log-odds score for CpG island being classified as slowly evolving.**

Scatter plot showing with a gray dot for each human CpG island the mean LECIF score (x-axis) and the log-odds score for classifying the CpG island as slowly evolving as opposed to quickly evolving (y-axis) from a previous study on primate CpG island sequence evolution[10]. In black circles the mean log-odds score for CpG islands binned by the LECIF score with ten equal-width bins is shown. These circles are connected with lines based on piecewise linear interpolation. One hundred thousand random human CpG islands annotated with the LECIF score were sampled to plot this scatter plot. PCC and SCC computed between the two scores across all CpG islands annotated with the LECIF score are shown in the bottom right. This illustrates that the LECIF score is positively correlated with the likelihood of a human CpG island being classified as slowly evolving as opposed to quickly evolving.

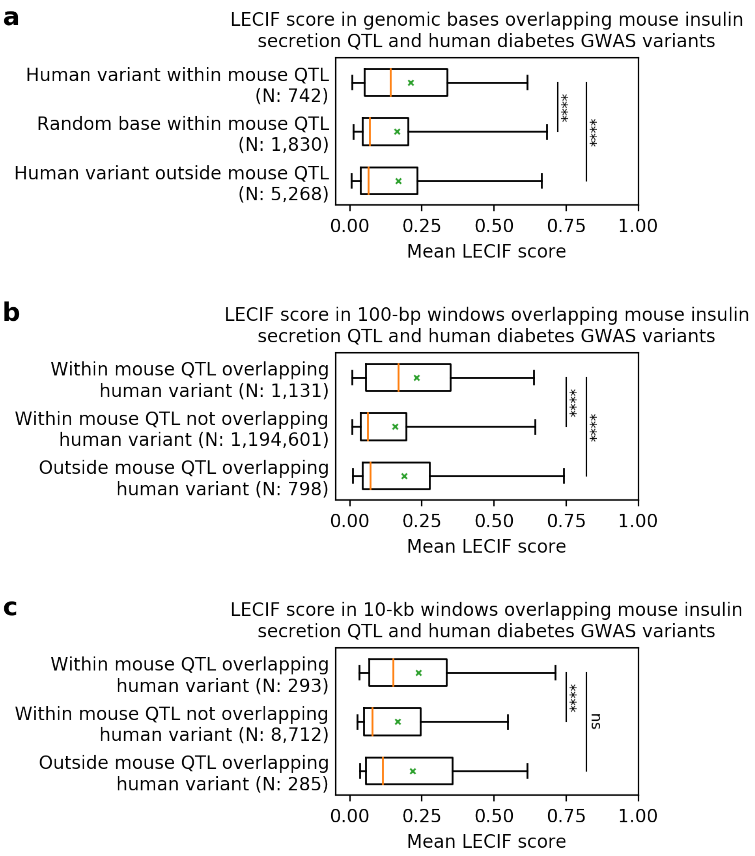


**Supplementary Figure 15. Distribution of mean LECIF score of human genomic windows overlapping mouse insulin secretion QTL and human diabetes GWAS variant.**

Similar to **Fig. 6c**, but showing the distribution of mean LECIF score identified as containing a human diabetes GWAS variant or overlapping a mapped mouse insulin secretion QTL or both[11] for non-overlapping **a.** 1-bp (genomic base), **b.** 100-bp, and **c.** 10-kb genomic windows, instead of the 1-kb windows shown in **Fig. 6c**. In each subplot, the top group refers to windows or bases that lie within the mouse QTL mapped to human and overlap or is the human GWAS variant. The middle group refers to windows or bases within the mouse QTL that do not overlap or is not the human GWAS variant. The bottom group refers to windows or bases randomly sampled from the human genome that lie outside the mouse QTL, but overlap or is the GWAS variant. Displayed after each label is the number of windows or bases corresponding to that group. Each distribution is represented by a boxplot with median (orange solid line), mean (green 'x'), 25$^{th}$ and 75$^{th}$ percentiles (box), and 5th and 95th percentiles (whisker). Any window with less than half of the bases annotated with the LECIF score was excluded. **** denotes p-value below 0.0001, and ns denotes p-value above 0.05 based on a Mann-Whitney U test.

**a** shows that human diabetes GWAS variants that overlap mouse insulin secretion QTL tend to score higher than the GWAS variants outside of the mouse QTL or bases that are not GWAS variants but within the mouse QTL. **b** and **c** show the result of **Fig. 6c,** that human genomic windows that overlap both mouse insulin secretion QTL and human diabetes GWAS variant tend to score higher than windows that overlap only one of them and that this result also holds for other window sizes.

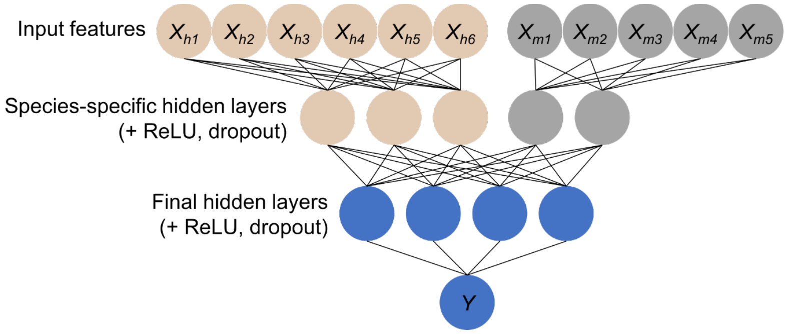


**Supplementary Figure 16. A schematic of a pseudo-Siamese neural network.**
A pseudo-Siamese neural network consists of two distinct sub-networks that do not share any weights[12]. The sub-network on the left (beige) takes in human feature vectors, $X_h$, and the sub-network on the right (grey) takes in mouse feature vectors, $X_m$. A final network (blue) takes in concatenated output vectors from the two sub-networks and generates the final prediction, $Y$. Each layer within a sub-network is followed by a rectified linear unit (ReLU) and dropout is used in the training[13]. We only show a small number of input features, layers, and neurons here. **Supplementary Table 3** lists the hyper-parameters that define this architecture.
